# A LINCS microenvironment perturbation resource for integrative assessment of ligand-mediated molecular and phenotypic responses

**DOI:** 10.1101/2021.08.06.455429

**Authors:** Sean M. Gross, Mark A. Dane, Rebecca L. Smith, Kaylyn Devlin, Ian McLean, Daniel Derrick, Caitlin Mills, Kartik Subramanian, Alexandra B. London, Denis Torre, Cemal Erdem, Nicholas Lyons, Ted Natoli, Sarah Pessa, Xiaodong Lu, James Mullahoo, Jonathan Li, Miriam Adam, Brook Wassie, Moqing Liu, David Kilburn, Tiera A. Liby, Elmar Bucher, Crystal Sanchez-Aguila, Kenneth Daily, Larsson Omberg, Yunguan Wang, Connor Jacobson, Clarence Yapp, Mirra Chung, Dusica Vidovic, Yiling Lu, Stephan Schurer, Albert Lee, Ajay Pillai, Aravind Subramanian, Malvina Papanastasiou, Ernest Fraenkel, Heidi S. Feiler, Gordon B. Mills, Jake Jaffe, Avi Ma’ayan, Marc R. Birtwistle, Peter K. Sorger, James E. Korkola, Joe W. Gray, Laura M. Heiser

**Affiliations:** Department of Biomedical Engineering, OHSU, Portland, OR, USA; Laboratory of Systems Pharmacology, Department of Systems Biology, Harvard Program in Therapeutic Science, Harvard Medical School, Boston, MA, USA; Department of Pharmacological Sciences, Mount Sinai Center for Bioinformatics, Icahn School of Medicine at Mount Sinai, New York, NY, USA; Department of Chemical and Biomolecular Engineering, Clemson University, Clemson, SC, USA; Broad Institute of MIT and Harvard, Cambridge, MA, USA; Department of Biological Engineering, Massachusetts Institute of Technology, Cambridge, MA, USA; Sage Bionetworks, Seattle, WA, USA; Sylvester Comprehensive Cancer Center, University of Miami, FL 33136, USA; Department of Molecular and Cellular Pharmacology, Miller School of Medicine, University of Miami, Miami, FL 33136, USA; Institute for Data Science & Computing, University of Miami, FL 33136, USA; Department of Genomic Medicine, Division of Cancer Medicine, The University of Texas MD Anderson Cancer Center, Houston, TX; Heart, Lung, and Blood Institute, National Institutes of Health, Bethesda, USA; Human Genome Research Institute, National Institutes of Health, Bethesda, USA; Department of Cancer, Developmental and Cell Biology, OHSU, Portland, OR, USA; Knight Cancer Institute, OHSU, Portland, OR, USA

**Keywords:** perturbation, phenotype, integrative analysis, module

## Abstract

The phenotype of a cell and its underlying molecular state is strongly influenced by extracellular signals, including growth factors, hormones, and extracellular matrix. While these signals are normally tightly controlled, their dysregulation leads to phenotypic and molecular states associated with diverse diseases. To develop a detailed understanding of the linkage between molecular and phenotypic changes, we generated a comprehensive dataset that catalogs the transcriptional, proteomic, epigenomic and phenotypic responses of MCF10A mammary epithelial cells after exposure to the ligands EGF, HGF, OSM, IFNG, TGFB and BMP2. Systematic assessment of the molecular and cellular phenotypes induced by these ligands comprise the LINCS Microenvironment (ME) perturbation dataset, which has been curated and made publicly available for community-wide analysis and development of novel computational methods (synapse.org/LINCS_MCF10A). In illustrative analyses, we demonstrate how this dataset can be used to discover functionally related molecular features linked to specific cellular phenotypes.

## INTRODUCTION

The function of cells and their organization into tissues is controlled by interactions between cell-intrinsic molecular networks and cell-extrinsic signals, and dysregulation of these signals is associated with various diseases^1^. Extracellular ligands activate cell surface receptors to modulate chromatin, RNA, and protein networks that induce changes in multiple cellular phenotypes including viability^2^, growth rate^3^, motility^4^, polarization and differentiation state^5^. Disease-specific studies—especially those focused on cancer—have concentrated on understanding phenotypes related to disease progression, resistance mechanisms, therapeutic vulnerabilities and molecular predictors of response^6–15^. Several canonical signaling pathways have been linked to distinct normal and disease-associated cellular phenotypes, including MAPK^16^, JAK/STAT^17^, WNT^18^, and TGFB^19^. However, a detailed mapping of the linkage between multi-modal molecular and phenotypic responses underlying cell state regulation, developmental processes and diverse diseases is lacking.

Two general approaches have been used to explore the role of extracellular signals in modulating cellular and molecular phenotypes. One approach involves systematic large-scale perturbation of panels of immortalized cell lines, which has yielded libraries of response signatures^6,8–11,13,20–22^. The other approach involves more focused assessment of phenotypic and molecular changes in more complex model systems, including engineered organoids^23,24^, flies^25^, worms^26,27^, fish^28^ and mice^29^. Together these studies indicate that comprehensive multi-omic assessment of perturbation responses is critical for gaining insights into molecular-phenotype relationships. Module analysis of multi-omic molecular data has proven a useful approach to identify co-regulated molecular features associated with normal^30–33^ and disease-associated^34^ phenotypes. Such data-driven approaches require comprehensive, systematically-generated datasets, and in recognition of this, multiple data generation and atlasing consortia have emerged over the past 20 years, including ENCODE^35^, TCGA^36^, GTEx^37^, and HubMap^38^.

The Library of Integrated Network-based Cellular Signatures (LINCS) consortium study presented here is a large-scale, cell line-based perturbation experiment designed to examine the molecular and phenotypic responses of normal cells to perturbations. Its uniqueness lies in the coordinated measurements of a large number of different cellular and molecular responses to biologically relevant ligands that, when studied together, can be used for systems-level analysis of microenvironmental responses. We focused on the well-characterized human mammary epithelial MCF10A cell line^39,40^, which has been extensively used to study tissue development^41^, migration^42,43^, and organoid formation^44,45^. The focus on a single cell line provided a controlled cell-intrinsic genetic context and also affords molecular and temporal density in experimental measurements. We studied responses to six ligands known to activate different canonical signaling pathways of biological and clinical relevance, enabling comparison of distinct molecular and phenotypic effects. These data are publicly available for community study at synapse.org/LINCS_MCF10A. The following sections describe and evaluate the information content of the LINCS ME perturbation dataset and present illustrative analyses showing how the dataset can be used to (a) elucidate molecular and cellular phenotypes that are influenced by the binding of specific ligands, (b) identify ligand-induced signatures that can be mined for biological insights, (c) discover candidate causal or functional relationships between molecular features with module analysis, and (d) identify molecular programs that control specific cellular phenotypes.

## RESULTS

### Approach to generate a LINCS ME perturbation dataset

Eight laboratories supported by the NIH LINCS program contributed to the creation and analysis of an MCF10A perturbation dataset to enable community study of the molecular mechanisms engaged by microenvironmental signals to modulate specific cellular phenotypes (**Fig. 1A**). **Figure 1B** shows the experimental and computational steps involved in the creation of the database. The process began with selection of ligands that strongly modulated phenotype. Both phenotypic and molecular responses to ligands were then measured over time and integrated computationally to identify the phenotypes and molecular modules engaged by each ligand. **Figure 1C** shows the experimental design in which multiple endpoints were measured at several time points after the introduction of ligands. The ligands and experimental assays are summarized in **Figure 1D**.

**Figure 1.**
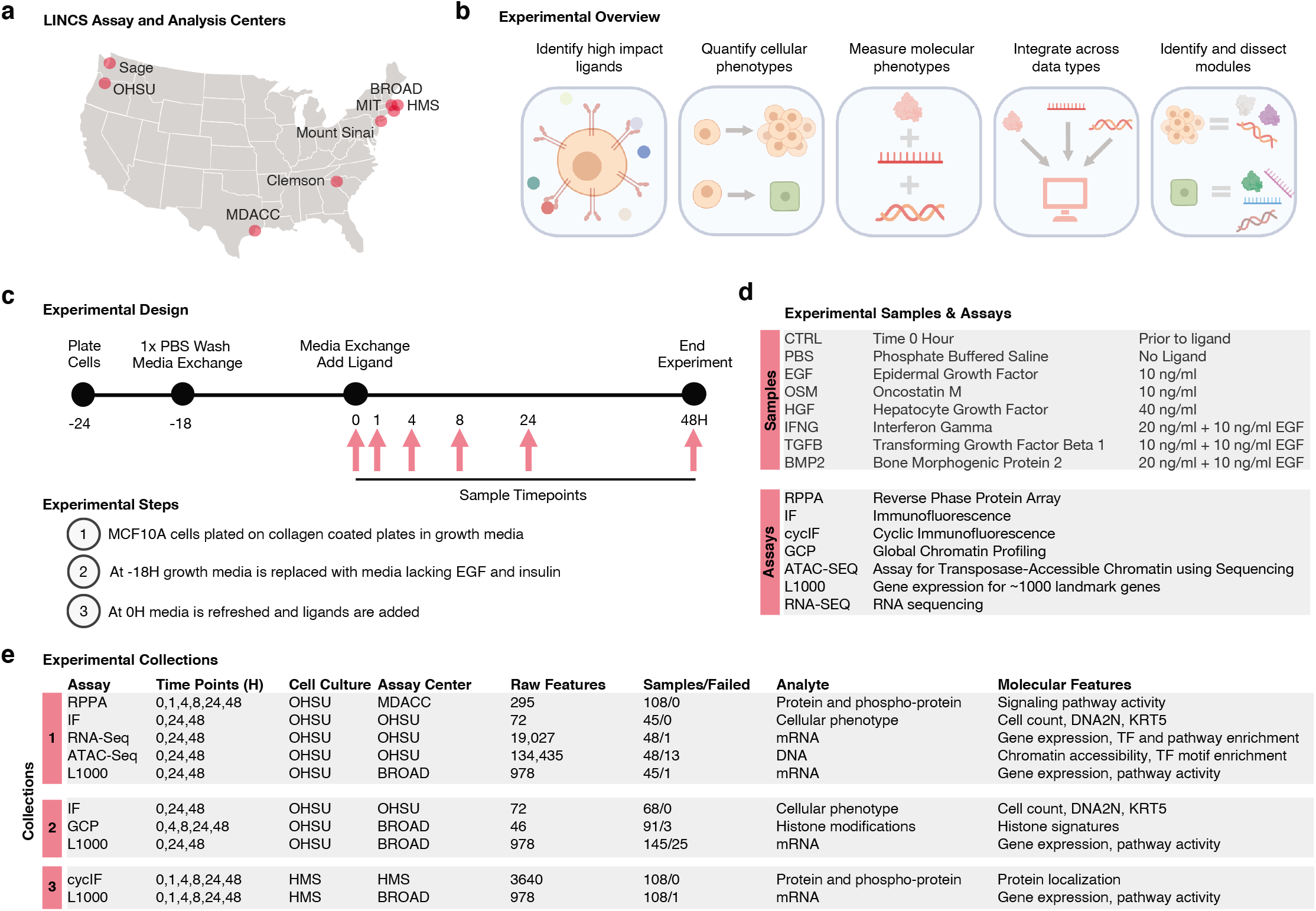
Overview of experimental approach to assess the impact of microenvironmental factors. (A) Map of LINCS data generation and analysis centers. (B) Schematic illustrating the experimental and analytical approaches to link molecular and cellular phenotypes. (C) Schematic of the experimental design, cell culture protocol, and sample harvest time points. (D) The experimental treatments, dosages, and assays deployed to generate the LINCS ME perturbation datasets. (E) Summary of the assays, timepoints, and features for the three experimental collections.

We selected six ligands based on the results of two high-throughput microenvironment microarray (MEMA) screens of 3024 combinations of 63 soluble ligands and 48 insoluble ECM proteins^46^; one screen with and another without EGF, a typical component of MCF10A growth medium^39^. We focused on collagen-1 as the insoluble ECM component and selected EGF, HGF, and OSM as ligands that increased growth in the absence of EGF; and BMP2, IFNG, TGFB as ligands that decreased growth in the presence of EGF (**Sup. Fig. 1A,B**.). These ligands target highly expressed receptors that are members of different canonical receptor classes (**Supp. Fig. 1C**). Dose-response experiments identified the ligand doses necessary to yield maximal changes in cell numbers (**Sup. Fig. 1D,E**). Inclusion of EGF in combination with BMP2, IFNG, and TGFB ensured sufficient cell numbers for molecular profiling.

The participating LINCS consortium laboratories performed systematic and large-scale analyses of epigenomic, transcriptomic, proteomic and phenotypic responses to each ligand at several time points during a 48H period after treatment (**Fig. 1B,D,E**). Cells for all analyses were grown and treated at OHSU and the treated cells or lysates were distributed to participating laboratories for analyses, except for those analyzed using cyclic immunofluorescence (cycIF)^47,48^. Cells for cycIF were grown and treated at HMS using cells, culture media and ligands supplied by one laboratory at OHSU to minimize experimental variation^49^ (**Fig. 1E**). For each assay, MCF10A cells were plated on collagen-1-coated cell culture dishes in their standard growth medium, which contains the growth factors EGF and insulin^39^. After attachment, the growth medium was replaced with medium lacking EGF and insulin, and cells were then treated with the ligand panel at optimized concentrations (**Figure 1D**).

Samples were collected before and after treatment over the 48H time period beginning with a time 0H sample (referred to as control: CTRL, **Fig. 1D**). Cellular responses were measured using live-cell imaging, four-color fluorescence imaging and cycIF^47,48^. Molecular responses were assessed for changes in protein expression with reverse phase protein arrays (RPPA)^50^; chromatin profiling using an Assay for Transposase-Accessible Chromatin using sequencing (ATACseq) and global chromatin profiling (GCP)^51^; RNA expression using RNAseq and the L1000^20^ transcriptomics panel designed to assess the levels of 1000 RNA transcripts selected to capture most of the variability in gene expression. Samples for the different assays were collected in three experimental collections comprised of at least three biological replicates each (**Fig. 1E**). Logistical and cost constraints resulted in some assays being applied to only a subset of time points. Rigorous quality assessment of all data led to the elimination of ~5% of samples (44/814). The resultant data and metadata are available at: synapse.org/LINCS_MCF10A.

### Overview of the ligand-induced cellular and molecular responses that comprise the LINCS ME perturbation dataset

#### Cellular responses

We quantified four-color immunofluorescence images from cells 24 and 48 hours after ligand treatment to assess cell clustering, cell density, shape, DNA content, and expression of proteins related to differentiation state (**Fig. 2A, Supp Table 1**). CycIF collected at all timepoints revealed additional changes in cell state and pathway activity. Consistent with our MEMA screen, HGF, OSM and EGF increased cell numbers and EdU incorporation (a measure of proliferation). BMP2 and TGFB significantly suppressed growth relative to the EGF condition; IFNG also reduced growth. (**Fig. 2C,D**). HGF, OSM, and IFNG+EGF upregulated KRT5 expression, a marker of basal differentiation state in mammary epithelial cells^52^ (**Fig. 2E**). OSM caused cells to form tight clusters (**Fig. 2F**). Lastly, TGFB+EGF induced evenly distributed cells with increased size, quantified as an increase in the distance to neighboring cells (**Fig. 2G**).

**Figure 2.**
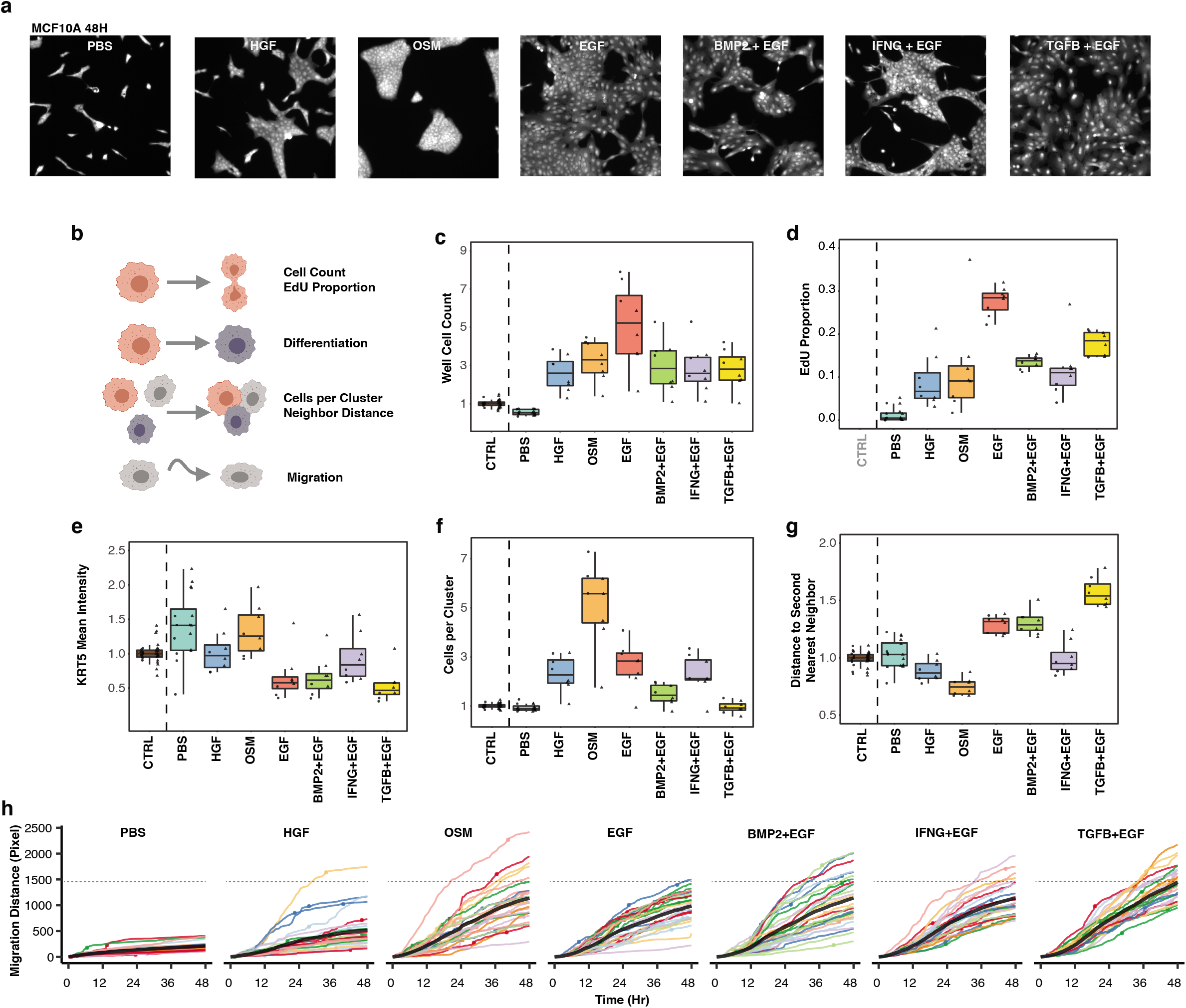
Ligand treatments induce diverse phenotypic responses. (A) Representative immunofluorescent images of ligand-induced cellular phenotypes at 48 hours. MCF10A cells were stained with Cell Mask to visualize cytoplasm. (B) Cartoon showing the image-based cellular phenotypes assessed from the immunofluorescence and live cell imaging assays. (C-G) Boxplots summarizing cellular phenotypes at time 0H (CTRL) and 48H after ligand addition from 8 biological replicates. Individual datapoints represent well-level means normalized to 0H. Circles are from collection 1 and triangles are from collection 2. Note that EdU positive proportion was not measured at 0H. (H) Accumulated cell migration (colored lines) from 0-48H for 25 cell lineages (individual cells and one of their progeny if they divided). Circles indicate mitotic events. The solid black lines indicate the population average; the dotted gray line shows the average TGFB+EGF induced migration at 48H, which was the treatment that induced the greatest increase in cell migration.

Analysis of live-cell images showed the emergence of each phenotype following ligand treatment (**Supp Movies**). OSM induced cells to undergo collective migration, a unique phenotype among the tested ligands. We quantified cell migration by tracking individual cells across the 48 hour time period and quantified migration as the total distance traversed by each cell lineage (**Fig. 2H**). In all ligand conditions, cell migration increased compared to the PBS condition, but to varying degrees: HGF-treated cells migrated the least while TGFB+EGF induced the greatest migration (Tukey’s HSD, p-value<.05). Together, the live cell imaging and migration analyses show the dynamic emergence of distinct phenotypic responses by each of the ligand treatments.

#### Molecular responses

The responses to ligands involved numerous features in each of the molecular datasets. Here we demonstrate some of our key observations through analysis of the RPPA proteomic dataset as an exemplar use-case. We assessed the modulation of canonical signaling proteins downstream from each ligand (**Fig. 3A**). These included: IRF1, a transcriptional target of STAT1 downstream of IFNG; pSTAT3, a signaling pathway component for OSM; and phosphorylation of MET, the receptor for HGF. PAI-1 provided an assessment of SMAD transcriptional activity, which is downstream of TGFB and BMP2. Additionally, phospho-HER2 provided a readout for conditions that contained EGF in the media. Each of these features were modulated as expected based on prior literature, validating the robustness of the dataset.

**Figure 3.**
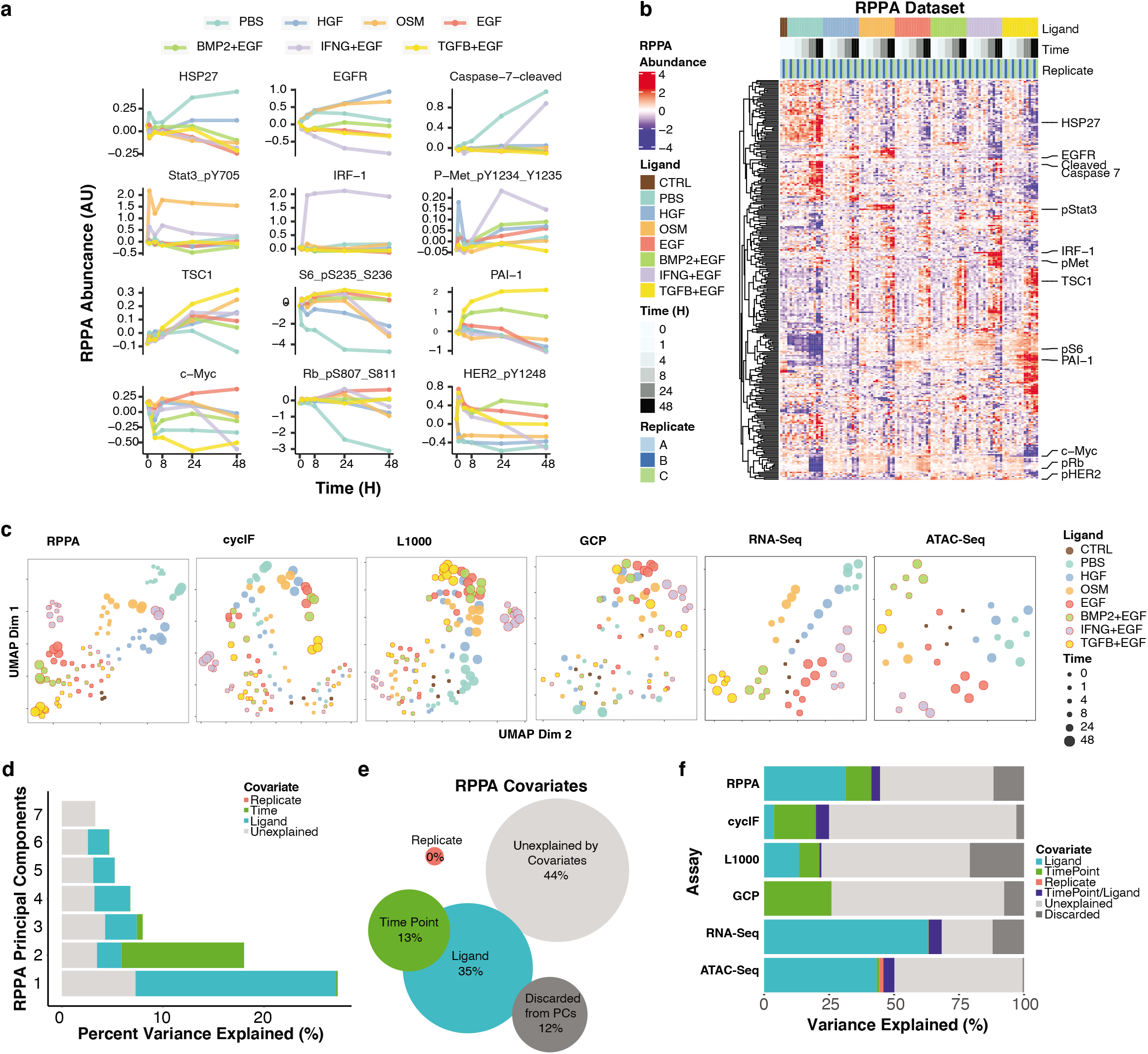
Six molecular assays reveal diverse dynamic responses to treatments. Line graphs show dynamic responses for 12 RPPA proteins under the different ligand treatments. (B) Heatmap of protein abundances as measured by RPPA. Rows represent abundance of 295 (phosphor)proteins and are median-centered and hierarchically clustered. Columns represent individual replicate samples, ordered by treatment and time. Callouts show the 12 proteins from panel A. (C) UMAPs for each of the six molecular assays. Each dot represents data from an individual sample and is the 2-dimensional embedding of all features measured in the assay. Color indicates ligand treatment and size indicates time point. (D) Plot of the first two principal components of RPPA assay. Variance in PC1 and PC2 is largely driven by ligand treatment and experimental timepoint, respectively. (E) MAVRIC analysis of RPPA covariates reveals the proportion of variance explained by sample replicate, experimental timepoint, and ligand treatment for each of the top seven principal components of the RPPA dataset. (F) Stacked bar graph shows a comparison of the information content contained within each molecular assay, as assessed by MAVRIC.

Unsupervised hierarchical clustering of the RPPA data set revealed dynamic changes in the protein landscape over time, with some responses shared by multiple ligands and others that were uniquely induced (**Fig. 3B**). All treatments that included EGF induced proteins related to growth factor signaling (*e.g*. pS6). The PBS condition, which lacks added growth factors, showed protein changes associated with reduced proliferation (*e.g*. decreased pRB) and induction of apoptosis (*e.g*. cleaved caspase 7), indicating that absence of growth factor signals strongly modulates phenotypic and molecular state.

To gain a high-level view of the six molecular assays, we performed Uniform Manifold Approximation and Projection (UMAP)^53^ dimensionality reduction for all ligand-induced responses (**Fig. 3C**). Most assays showed ligand-specific effects, as observed by samples from the same ligand treatment tending to group near one another. In addition, most datasets showed evolution over time from the starting state to another distinct state, captured by early time points clustering near the center of the UMAP and later time points for each ligand appearing in different UMAP regions.

#### Assessment of assay variance

We applied the Measuring Association between VaRIance and Covariates (MAVRIC) method to systematically assess the fractional variance explained by each experimental covariate of ligand, time, and replicate^54,55^. In brief, we first performed principal component analysis (PCA) to reduce the dimensionality of each data set while preserving the variability. Next, we quantified the total variance explained by each covariate (ligand, time, replicate) by summing the weighted variances of all statistically significant principal components (PCs). For example, in the RPPA dataset, the signal in the first PC was dominated by ligand while the second PC was dominated by time point (**Fig. 3D**). Summing across all significant PCs from the RPPA dataset revealed that 35% of the variance could be attributed to ligand and 13% to time point (**Fig. 3E**). Variance explained by multiple co-variates is represented by overlap in the Venn diagram. Overall, 44% of the variance in the RPPA dataset could not be explained by one of these factors, suggesting signal in the data attributable to other factors, such as changes shared by multiple ligands. Similarly, all other assays carried signal attributable to ligand treatment, although to varying degrees: RNAseq (63.1%) and ATACseq (43.3%) contained the greatest ligand-associated signal while GCP (0.1%) contained the least (**Fig. 3F**). Datasets with both early and late time points (RPPA, GCP, cycIF) carried signal attributable to time. There was limited variation attributable to replicates across all assays, indicating modest biological and technical variation.

### Identification and analysis of ligand-induced molecular signatures

Here we present a systematic assessment of molecular signatures induced by each ligand and provide examples of how these signatures can be analyzed and mined. Specifically, we focus on IFNG+EGF to examine the temporal evolution of responses across modalities and to identify novel immune-related molecular features.

#### Identification of ligand-induced signatures

To create molecular signatures of ligand responses, we identified features from each of the 6 data types that were differentially expressed at 24H and 48H timepoints relative to the CTRL sample (q-value < 0.01, |logFC| ≥ 1.5) (**Fig. 4A**). Features were classified as ligand-unique if they were modulated by a single ligand or shared if they were induced by more than one treatment (**Supp Tables 2,3**). All treatments induced both ligand-unique and shared molecular responses. IFNG+EGF, TGFB+EGF and OSM induced the greatest molecular changes as measured by the combination of RNAseq, ATACseq, GCP, cycIF and RPPA, indicating robust shifts in molecular state. In contrast, EGF, HGF and BMP2+EGF showed more modest effects, consistent with maintenance of MCF10A cells in a pre-treated state. Cross-correlation analysis of the molecular responses revealed that 24H and 48H responses were strongly correlated for each ligand and that responses to ligands from related families were more similar to one another than to other family classes (BMP2/TGFB, OSM/IFNG, EGF/HGF) (**Fig. 4B**, **Supp Table 4**).

**Figure 4.**
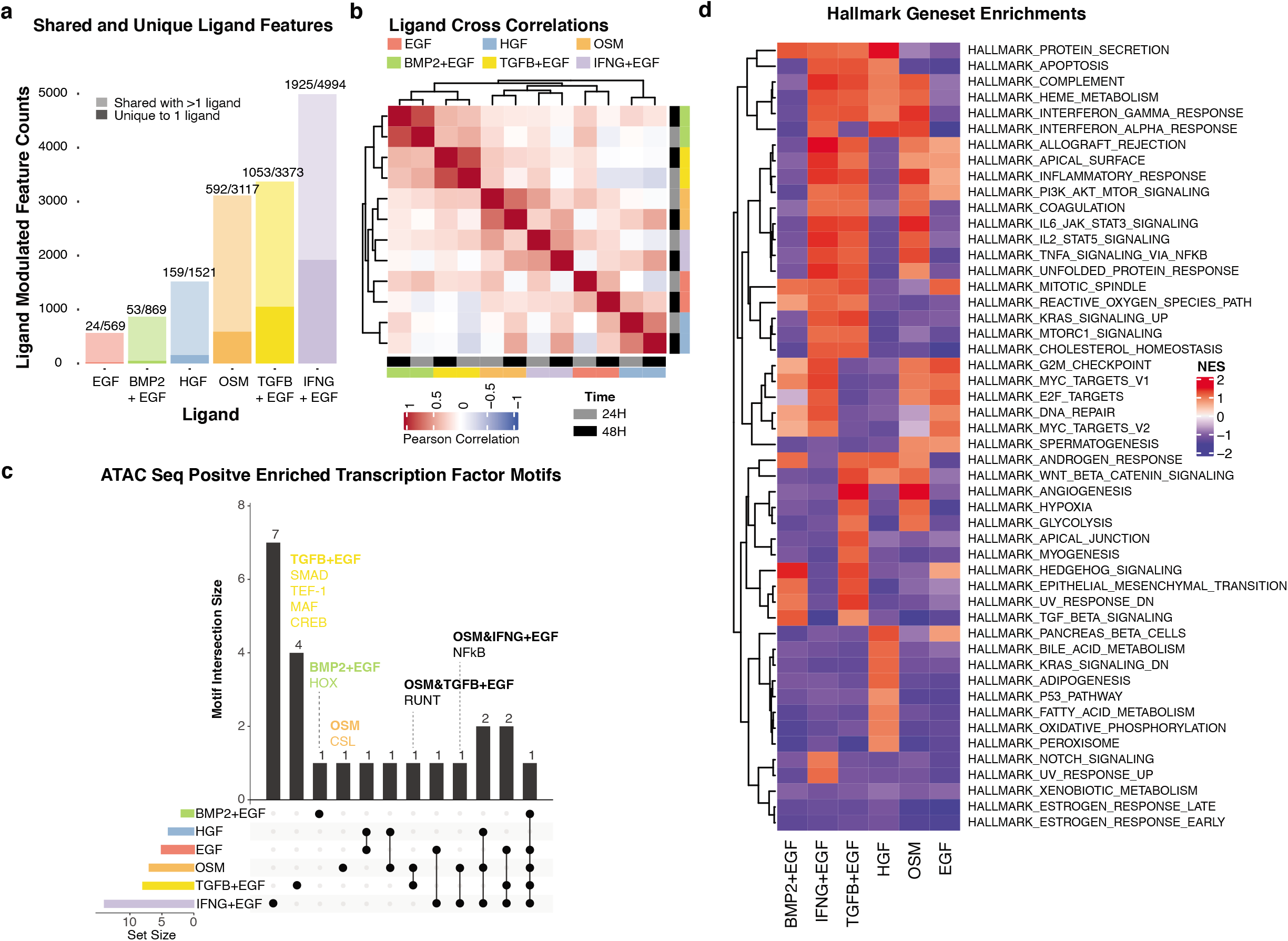
Assessment of ligand-induced molecular change. (A) Barplot showing the number of features significantly modulated by each ligand treatment. Shading indicates whether induced features are unique to a particular treatment (dark) or induced by multiple treatments (light). Numbers above bars indicate the number of features uniquely induced over the total number of features induced. (B) Heatmap showing pairwise correlations between molecular features induced by each ligand. Ligand responses from similar families are more highly correlated than those from unrelated families. (C) UpSet plot showing overlaps of induced ATACseq transcription factor motifs among ligand treatments. Column heights represent the number of transcription factor motifs induced by the ligand(s) indicated with filled dots. (D) Hallmark Geneset enrichment scores computed from RNAseq data.

Motivated by our observation from the MAVRIC analysis that the ATACseq and RNAseq datasets carried the strongest ligand signals, we more deeply interrogated these responses. We analyzed ATACseq transcription factor binding motif enrichment, a measure of transcription factor activity, and found that IFNG+EGF and TGFB+EGF induced the greatest number of enriched motifs. For example, TGFB+EGF induced SMAG, TEF-1, MAF and CREB motifs, while TGFB+EGF and OSM both induced changes in RUNT. (**Fig. 4C**). Gene set enrichment (GSEA) analysis^56^ of the RNAseq dataset revealed a unique complement of gene programs associated with response to each ligand treatment (**Fig. 4D**, **Supp Table 5**). To identify targeted inhibitors that induce similar molecular mechanisms, we compared our ligand signatures against the LINCS L1000 database^57^ of drug and other chemical response signatures (Fisher exact test, q-value<0.2). The ligand panel activated many of the same molecular signatures as small molecule inhibitors profiled in the L1000 database, indicating that many small molecules induce similar molecular responses as the ligands. This analysis also suggests that environmental signals may modulate therapeutic response (**Supp Fig 2, Supp Table 6**).

#### Identification of novel molecular features induced by IFNG

We analyzed responses to IFNG+EGF to illustrate how the LINCS ME perturbation dataset can be used to study the molecular mechanisms associated with ligand responses across time. IFNG is a soluble cytokine secreted by cells of both the innate and adaptive immune systems and has become increasingly scrutinized, owing to interest in understanding the role of the immune system in diverse pathophysiologies^58^ as well as cancer immunotherapies. IFNG+EGF treatment induced dynamic changes in canonical IFNG signaling molecules measured across assays, including: rapid nuclear translocation of STAT1 and induction of IRF1, followed by upregulation of PDL1 at the membrane and associated epigenetic changes (**Supp Fig. 3A-F**). These findings indicate that the LINCS ME perturbation dataset enables the encoding of a stimulus to be traced across time and molecular modalities.

We observed that 66/202 Pathcards Reactome IFNG superpathway features^59^ were among the most strongly induced by IFNG+EGF treatment, indicating the induction of multiple known signaling responses (**Supp Fig. 3G**). To gain deeper insight into the ability of IFNG to influence both adaptive and innate immune responses through altering cytokine production by tumor cells, we compared the MCF10A IFNG+EGF signature, the IFNG superpathway, and a curated cytokine gene list^60^. This comparison identified 15 cytokines not already included in the IFNG superpathway, suggesting additional cytokines produced by tumor cells in response to IFNG that may interact with various immune cell subsets, including: CSF1^61,62^, IL15^63^, IL12A^64^, CCL2^65^, and CXCL2^66^. This demonstrates how the LINCS ME dataset can be mined to gain novel biological insights into immune-related signaling and to prioritize molecular features for future study.

### Discovery of candidate functional relationships between molecular features

We reasoned that the patterns of robust multi-omic molecular changes induced across the panel of ligands could be analyzed together to discover coordinately regulated molecular programs. Importantly, our use of multiple ligands that perturb cells along various phenotypic and molecular axes enabled distinct molecular programs to be disentangled. Below we summarize our assessment of the relationships between different modalities, our approach to identify coordinately regulated biological modules, and also illustrate the utility of the modules to provide insights into the molecular programs active across diverse tissues.

#### Identification of coordinately regulated modules

We assessed coordinated responses in the RPPA, RNAseq, and ATACseq datasets by molecular cognates across datasets (*e.g*. Cyclin B1 in RPPA and CCNB1 in RNAseq) and found broad concordance, indicating conserved responses across molecular modalities (**Supp Fig. 4**). We next used a systematic approach to identify modules comprised of coordinately regulated molecular features measured in the different assays. Specifically, we examined all molecular features that were induced by at least one ligand (see **Fig. 4A**) and then scaled each assay dataset with an rrscale, which is a transformation that normalizes feature distributions, removes outliers, and z-scales feature values^67^ (**Supp Fig. 5**). Next, we clustered features with partitioning around medoids (PAM) followed by gap statistic analysis to identify the optimal number of clusters. This yielded 18 molecular modules; highly correlated modules were combined to yield a final set of 14 molecular modules for interpretation (**Supp Fig. 6A-C**).

Each module represents a unique complement of co-regulated proteomic, transcriptional, and chromatin features (**Fig. 5A**). Features from each assay were well distributed across modules, with RPPA and RNAseq features represented in all modules; assay features not distributed across all modules were those with less comprehensive coverage of diverse biological pathways, such as cycIF and GCP (**Supp Fig. 6D, Supp Table 7**). Each module showed distinct modulation patterns across the ligands; most modules were induced by more than one ligand while a few were ligand-specific, consistent with the findings in **Figure 4**. Reactome pathway enrichment analysis demonstrated that each module induced an array of transcriptional programs (**Fig. 5B Supp Table 8**). Transcription Factor enrichment via ChEA3^68^ identified key molecular drivers associated with these gene programs (**Fig. 5C**, **Supp Table 9**).

**Figure 5.**
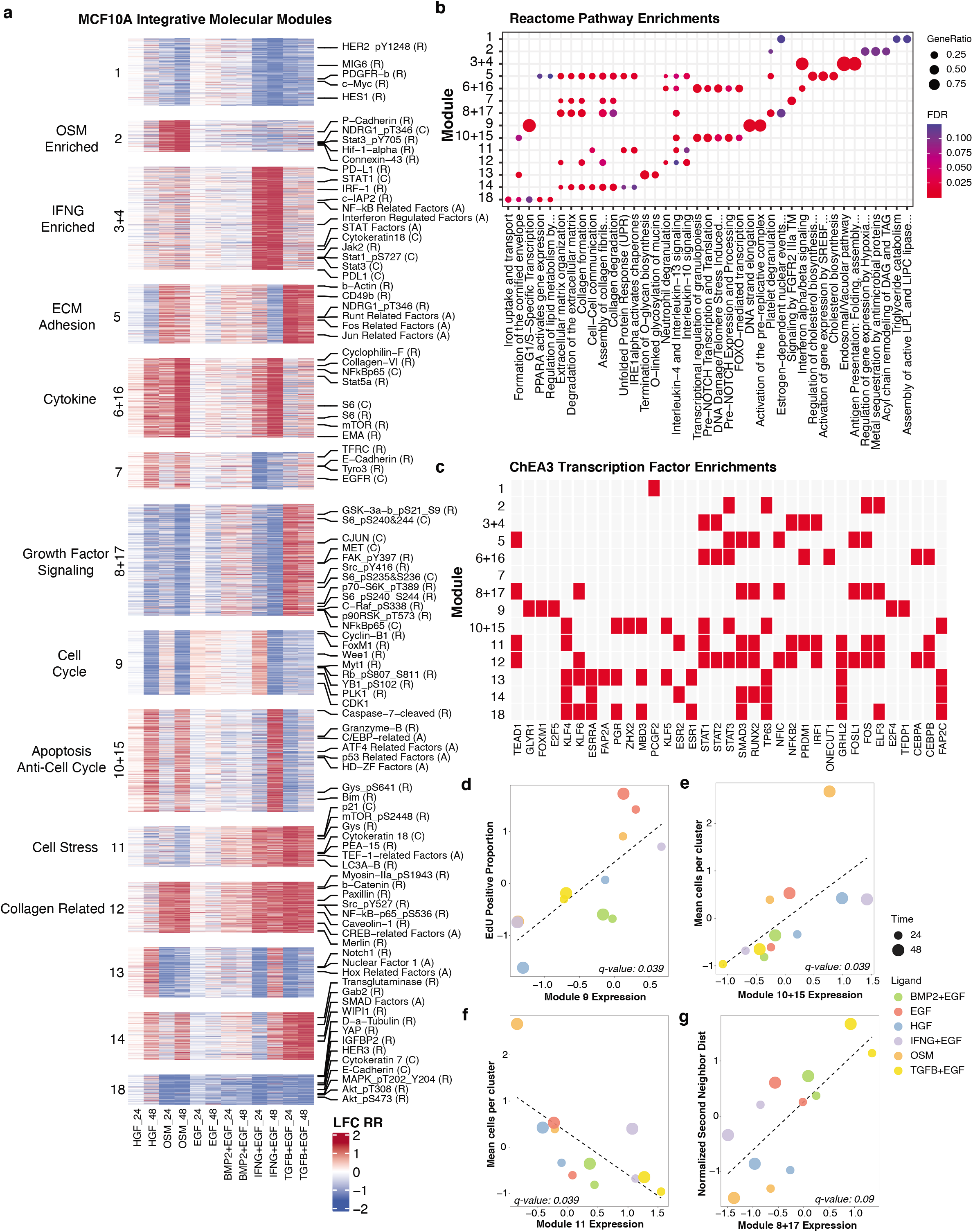
Integrated analysis identifies co-regulated molecular modules. (A) Heatmap showing the 14 integrative molecular modules for each ligand at 24H and 48H. Features are grouped by cluster. Biological interpretation for modules is indicated on the left; feature callouts for RPPA (R), cycIF (C), ATACseq (A) are shown to the right. (B) Bubble plot shows the top enriched Reactome pathways in each module, computed from RNAseq features. Dot size indicates the gene ratio; dot color indicates FDR value. (C) Heatmap shows results of ChEA3 transcription factor enrichment analysis computed for the RNAseq features in each module. Red indicates the top 5 TFs (by p-value) per module that met a p-value threshold of 0.05. (D-G) Scatterplots show the relationships between module activity and quantitative phenotypic responses for selected pairs. Dot color indicates the ligand treatment and dot size indicates the time point. The black dotted line shows the linear fit, and the q-value of the fit is shown at the bottom of the plot.

#### Assessment of molecular modules across diverse tissues

Elucidating the molecular programs operable across different tissue types is critical for understanding normal organ development and function, and also for identifying molecular programs that may go awry in the case of disease. We assessed RNA expression of the 14 integrated modules in the GTEx normal tissue dataset^37^ to identify molecular programs that may be most active in particular tissue types (**Supp Fig. 7, Supp Table 10**). We observed tissue-specific activation of the modules. For example, Module 6+16, which was enriched for immune-related programs, was highly expressed in multiple brain regions, consistent with the importance of immune signaling in brain function and neuroplasticity^69^. Module 10+15 included gene expression programs related to transcription, translation, and senescence and was highly expressed in GTEx pancreas samples. Supporting this, RNA processing has recently emerged as an important molecular function in the regulation of pancreatic beta-cells in normal and diabetic conditions^70,71^. Module 5 was enriched in extracellular matrix organization and collagen formation pathways and proteins associated with cell adhesion (CD49b (ITGA2)). This module was highly expressed in artery samples, consistent with the observation that the arterial wall produces a rich and complex extracellular matrix that defines the mechanical properties of the vessel^72,73^. Additional features included in each of these modules may further shed light on their roles in normal and diseased processes in different tissues.

### Investigation of the relationship between molecular modules and cellular phenotype

Elucidation of the molecular mechanisms that control cellular phenotype remains a difficult problem in systems biology. We illustrate here how the LINCS ME perturbation dataset can be analyzed to gain insights into mechanisms of phenotype control by linking cellular and molecular responses. We present two examples: a data-driven discovery of associations between quantitative phenotypic responses and module activity, followed by a detailed analysis of Module 2 to uncover molecular features associated with the unique cell clustering and collective motility phenotype induced by OSM.

#### Data-driven discovery of phenotype-module associations

We performed correlation analysis to identify molecular modules that were significantly associated with features measured in imaging-based assays (**Fig. 5D-G**). We found that Module 8+17 was positively correlated with the phenotypic response ‘Normalized Second Neighbor Distance’, a metric that reflects both cell size and cell-cell spatial organization (**Fig. 5G**, p-value = 0.005). Several features of this module suggest molecular correlates of this phenotypic response, including pathway enrichments in ECM-related programs and multiple phosphorylated growth factor signaling proteins. Additionally, the transcription factor RUNX2, which was enriched in this module, has been implicated in modulating cell morphology and cell spreading^74^. Finally, several GCP features suggest post-translational chromatin modifications associated with cell morphology changes.

We also identified a specific and robust correlation between Module 9 expression and the fraction of EdU positive cells (**Fig. 5D**, p-value = 0.001). To explore the putative regulatory components of Module 9, we annotated genes that code for transcription factors, kinases, noncoding RNA, and epigenetic regulators (**Fig. 6A, Supp Table 11**). This analysis revealed a suite of factors previously shown to play key roles in regulating cell cycle progression, including the transcription factors: E2F1, FOXM1, MYB, and TFDP1; and the kinases: AURKA, CDK1, PLK1, and BUB1. Module 9 RPPA features cyclin B, Wee1 and phosphorylated RB are canonical cell cycle proteins that showed temporal dynamics consistent with changes in proliferation, as well as lesser linked features including FOSL1 ^75–77^ and PASK^78,79^. (**Fig. 6B**). ChEA3 transcription factor enrichment^68^ identified multiple cell cycle-associated transcription factors including FOXM1, TFDP1 and E2F isoforms (**Fig. 6C**). The most significantly enriched Reactome pathways were cell cycle, DNA replication, and DNA repair (**Fig. 6D**). We analyzed the top 5 sub-pathways within each of these Reactome pathways and found the highest enrichment for G1/S specific transcription, PCNA-dependent base excision repair, and unwinding of DNA (**Fig. 6E**). Additionally, Module 9 included 86% (37/43) of the genes in a functionally-annotated G1/S gene set^80^, with expression patterns consistent with changes in EdU incorporation (**Fig. 6F**). There is also evidence for DNA damage and potentially for replication stress in the induction base-excision repair, the G2M checkpoint and activation of DNA damage checkpoint associated kinases. In sum, Module 9 contains cell cycle-associated molecular features from multiple modalities.

**Figure 6.**
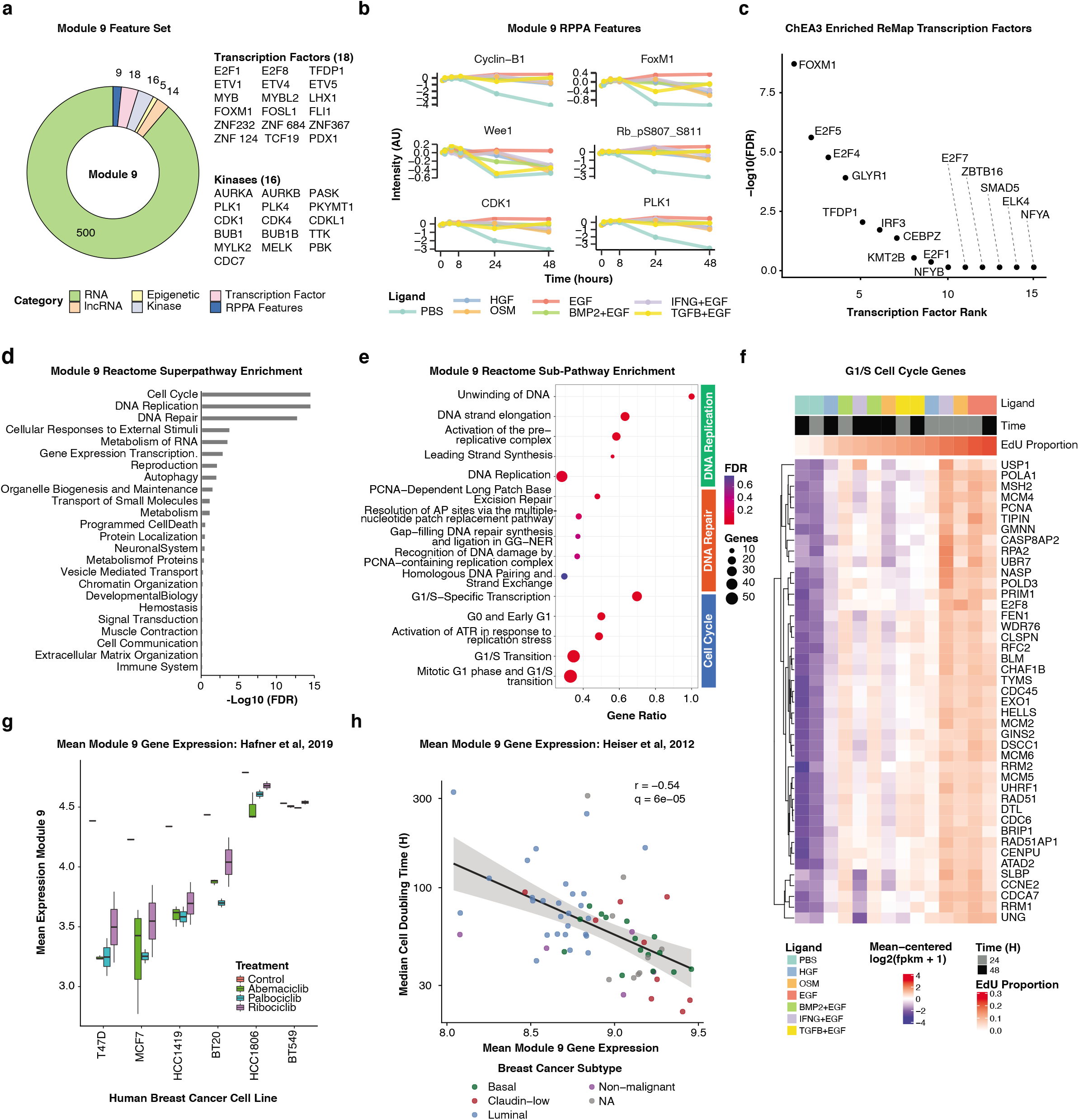
Module 9 is associated with cell cycle progression. (A) Donut plot showing distribution of Module 9 features across assays. Transcription factors and kinases in the RNA gene set are called out to the right of the plot. (B) Line plot showing 6 of the Module 9 RPPA features. (C) Plot of the top 10 most significantly enriched transcription factors inferred from the Module 9 RNA gene set. (D) Bar plot showing the enrichment of Reactome superpathways from the Module 9 RNA gene set. (E) Bubble plot showing the top 5 enriched Reactome subpathways from the Reactome Cell Cycle, DNA Repair, and DNA Replication superpathways. Dot color indicates q-value; dot size indicates the number of genes in Module 9 that are found in each gene set. (F) Heat map showing the expression of the Seurat G1/S cell cycle gene set, sorted based on the EdU positive proportion. (G) Plot of mean Module 9 gene expression for a panel of breast cancer cell lines treated with three CDK4/6 inhibitors for 24H or an untreated control. Data from Hafner, et al 2019. (H) Dot plot of mean Module 9 gene expression from 65 human breast cancer cell lines graphed against their mean doubling time. Cell lines are colored based on their breast cancer subtype classification. The line indicates the linear fit across all cell lines. Data from Heiser, et al 2012.

To test if the link between Module 9 and cell cycle control generalized beyond MCF10A cells, we analyzed two publicly available independently generated breast cancer cell line data sets. First, we quantified mean Module 9 gene expression scores from 7 breast cancer cell lines treated for 24 hours with a panel of CDK4/6 inhibitors^81^. As expected, this showed robust downregulation of Module 9 in response to each of the three CDK4/6 inhibitors in the five sensitive cell lines, while the two resistant cell lines showed only modest changes in Module 9 expression (t-test, p-value < 0.0001, **Fig. 6G**). In a second analysis, we compared Module 9 expression for a panel of 65 breast cancer cell lines^10^ against cell doubling time, which revealed a significant correlation, consistent with the interpretation that Module 9 is functionally associated with the cell cycle (**Fig. 6H**, Pearson R = −0.54). All together, these analyses indicate that our data-driven approach to module detection can identify coordinately regulated molecular features associated with quantitative phenotypic responses and that these findings generalize to independent data sets.

#### Examination of module activity to elucidate the molecular basis of ligand-induced phenotypic responses

In our final analysis, we illustrate how the modules can be examined to provide insights into the molecular basis of complex phenotypic responses. Here, we focused on OSM, a member of the IL6 cytokine family implicated in immune function, developmental processes, and tissue remodeling^82^. OSM stimulated proliferation and was the only ligand in our panel that induced collective migration, a complex phenotype in which individual cells form tight clusters that undergo migration (**Fig 7A**, **Supp Movies**).

**Figure 7.**
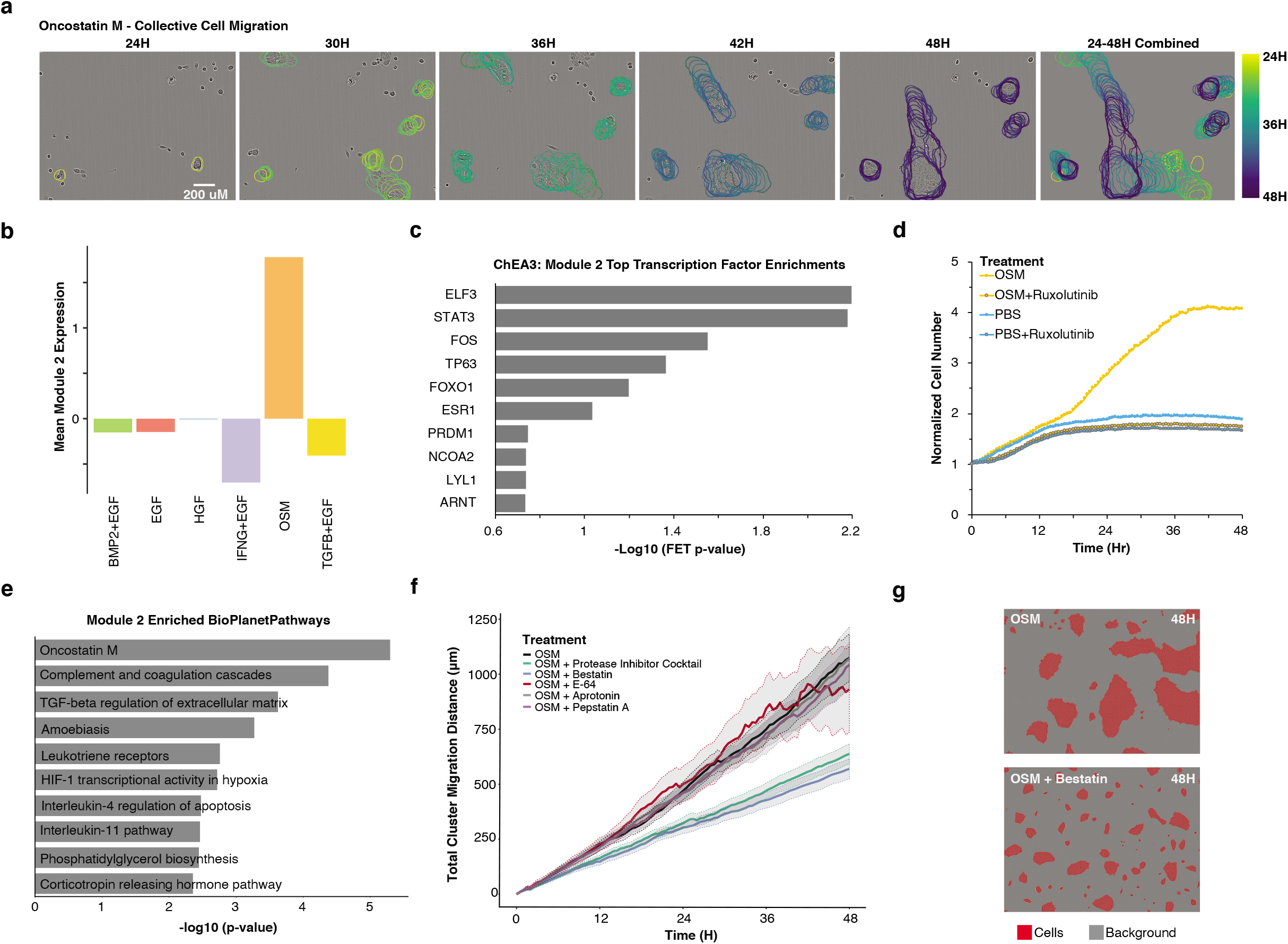
Analysis of molecular modules identifies functional relationships between molecular and phenotypic responses to OSM. (A) OSM induces the formation of cell clusters that undergo collective migration and merge to form large clusters. Representative tracks of OSM-induced cluster migration are shown from 24 hours to 48 hours after OSM treatment. Cluster outlines are colored by experimental timepoint. All images are set to the same scale. (B) Barplot shows the mean Module 2 expression for the six ligand treatments. (C) Barplot showing the top 10 enriched transcription factors inferred for the Module 2 genes in Chea3. (D) The JAK/STAT inhibitor Ruxolitinib inhibits cell growth in the presence of OSM. Line graph shows the relative number of cells across time. PBS (phosphate buffered saline) treatment serves as a control. (E) Barplot of the top 10 enriched pathways in Bioplanet using the module 2 RNAseq gene set (F) OSM-induced collective migration is mediated by protease activity. Line graph shows the accumulated cluster migration distance after OSM +/-a protease inhibitor cocktail and its individual components including bestatin, E-64, aprotonin, and pepstatin A. Solid lines show the population average and gray shaded regions indicate 95% confidence intervals of the mean distance travelled at each timepoint. (G) False color phase contrast images at 48H show that bestatin inhibits the formation of large cell clusters in the presence of OSM. Cells have been colored red and the background has been colored gray.

To gain insight into the molecular features underlying this unique phenotype, we focused on modules that were strongly induced by OSM, including Modules 2, 6+16 and 12 (**Supp Fig. 6**).

Features in Module 2 were of particular interest, as this module was selectively induced by OSM (**Fig. 7B**). Module 2 includes RPPA features pSTAT3, P-Cadherin, Connexin-43, and Hif-1-alpha as well as transcription factor enrichment in ELF3, STAT3, TP63, and FOS (**Fig. 7C, Supp Table 7**). P-Cadherin and Connexin-43 are intriguing, as they are implicated in the cell adhesion contacts required for mediating the observed clustering phenotype^83,84^. Based on the coordinated changes in STAT3 across modalities, we tested the functional importance of this axis with Ruxolitinib, a JAK/STAT inhibitor. We found that addition of Ruxolitinib in the presence of OSM strongly inhibited both the growth of cells and cell migration, confirming the importance of JAK/STAT signaling in mediating responses to OSM (**Fig. 7D, Supp Movies**).

To probe more deeply into the Module 2 RNAseq features, we tested for enriched pathways using BioPlanet^85^ (**Fig. 7E**). The top pathway hit in this analysis was ‘OSM’, which serves as a validation of the module approach. The second hit was ‘complement and coagulation cascades’, two linked processes driven by a series of proteases to stimulate innate immunity and blood clotting^86^. This suggested that protease activity may be critical for mediating OSM-induced cluster migration. To examine the role that proteases play in cluster migration, we treated MCF10A cells with OSM in the presence of a cocktail of five protease inhibitors, and found reduced cluster migration indicating the importance of protease activity in this phenotype (**Fig. 7F**). We next tested individual components of the protease cocktail and found limited effects of aprotinin, E-64, and pepstatin A. However, with bestatin, an aminopeptidase inhibitor, we observed formation of cell clusters but a failure of these clusters to migrate and merge together (**Fig. 7G**). Thus, these functional studies developed from the module analysis implicate aminopeptidase activity as a critical mediator of OSM-induced collective cell motility in MCF10A cells. Overall, our approach to leverage responses to multiple perturbations enabled identification of molecular programs associated with complex phenotypic responses including cluster migration and cell proliferation.

## DISCUSSION

Here we leveraged the LINCS Consortium framework to systematically quantify the phenotypic and molecular responses of MCF10A mammary epithelial cells after treatment with a diverse panel of ligands. Analysis of this dataset revealed robust molecular and phenotypic responses and enabled identification of ligand-specific signatures, integrated molecular modules, and linkage of phenotypic and molecular responses. These data support the idea that deeply examining a single model system subjected to a range of perturbations with measurements across multiple modalities is crucial to understanding complex biological phenomena.

The robust, multimodal dataset enabled a range of computational analyses. For instance, the coordinated use of a diverse panel of molecular assays facilitated comparisons of the information carried by each assay and revealed that RNAseq and ATACseq assays had the greatest ligand-associated signal. Differences in information content between assays may be due to: intrinsic differences in molecular modalities, the signal available in a particular assay, or differences in the number and diversity of biologically meaningful features in each assay. These findings suggest that comprehensive assays such as RNAseq are well-suited for discovery-based screens or experiments that examine large panels of perturbagens, whereas targeted assays such as cycIF—which can be adapted through inclusion of different biomarkers—would be expected to excel in focused hypothesis-driven studies^47,48^.

In our integrated analysis, we joined epigenomic, transcriptional and proteomic changes into coregulated modules. Critical for this analysis was the use of ligands that stimulate diverse and partially overlapping pathways, as this enabled identification of molecular features that were subtly and variably induced by multiple ligands. We analyzed the modules to identify linkages between molecular features and phenotypic responses. For instance, we identified a set of coregulated molecular features strongly associated with cell cycle, including both canonical transcriptional factors, pathways, and proteins as well as features that have been implicated but not confirmed in cell cycle regulation, such as PASK^78,79^. These analyses demonstrate how the LINCS ME Perturbation dataset can be used to formulate specific testable hypotheses that could be explored in future experimental studies. Some modules were semi-correlated and contained similar biological programs, as indicated by enrichment of shared pathways and TF programs. Alternate methods to identify modules that permit partial membership of individual features may allow a more nuanced identification of the relationship between molecular features^87^. Overall, this tightly controlled framework allowed molecular signals to be disentangled and associated with quantitated phenotypic responses.

Our live-cell imaging studies revealed the induction of phenotypic responses in response to ligand perturbation. In particular, OSM uniquely induced MCF10A cells to form tight cell clusters that underwent collective migration. We used our module analysis to explore the molecular basis of this complex phenotypic response and examined modules that were uniquely induced by OSM. Experimental validation identified functional links between OSM-induced molecular and phenotypic responses: protease activity was required for collective cell migration while STAT activation was required for proliferation. Our findings add to the substantial literature that implicates proteases in modulating interactions between cellular and extracellular signals^88^. Future studies that examine the role of other Module 2 features will be needed for a complete understanding of the molecular basis of OSM-induced collective migration. Finally, additional complex phenotypic responses could be investigated by growing MCF10A cells as 3D organoids^45^.

Together, our findings indicate that this LINCS ME perturbation dataset will serve as a robust and valuable resource for community-wide analysis and exploration. This resource can be utilized by the broader community to gain deeper insights into biological processes such as the molecular basis of different phenotypes, the molecular and phenotypic impact of particular ligands, and how particular molecular features are modulated by perturbation. Additionally, these data can serve as a resource for computational scientists to examine relationships between different molecular modalities, to develop methods for identifying molecular networks, or to elucidate the temporal relationships between different types of molecular changes. We also envision expansion of the dataset to include additional molecular measurements (*e.g*. single-cell RNAseq, single-cell ATACseq, and single-cell proteomics) and perturbation with different ligand combinations.

## METHODS

### Cell Culture Methods

#### Cell culture

To decrease heterogeneity MCF10A cells were frozen in a single batch at the MD Anderson Cancer Center and used by both OHSU and HMS from the frozen batch with limited passaging. Cell identity was confirmed by short tandem repeat (STR) profiling, and cells tested negative for mycoplasma. Cells were cultured in growth media (GM) composed of DMEM/F12 (Invitrogen #11330-032), 5% horse serum (Sigma #H1138), 20 ng/ml EGF (R&D Systems #236-EG), 0.5 μg/ml hydrocortisone (Sigma #H-4001), 100 ng/ml cholera toxin (Sigma #C8052), 10 μg/ml insulin (Sigma #I9278), and 1% Pen/Strep (Invitrogen #15070-063). For ligand treatments a growth factor free media was used, experimental media (EM), that was composed of DMEM/F12, 5% horse serum, 0.5 μg/ml hydrocortisone (Sigma #H-4001), 100 ng/ml cholera toxin (Sigma #C8052), and 1% Pen/Strep (Invitrogen #15070-063).

Prior to experiments, MCF10A cells were grown to 50-80% confluence in GM and detached using 0.05% trypsin-EDTA (Thermo Fisher Scientific 25300-054). Following detachment, 75,000 cells were seeded into collagen-1 (Cultrex #3442-050-01) coated 8-well plates (Thermo Fisher Scientific 267062) in GM. Approximately 7 hours after seeding, cells were gently washed with PBS and EM was added. Following 18 hours of incubation in EM, cells were treated with ligand in fresh EM media as follows: 10 ng/ml EGF (R&D Systems #236-EG), 40 ng/ml HGF (R&D Systems #294-HG), 10 ng/ml OSM (R&D Systems #8475-OM), 20 ng/ml BMP2 (R&D Systems #355-BM) + 10 ng/ml EGF, 20 ng/ml IFNY (R&D Systems #258-IF) + 10 ng/ml EGF, 10 ng/ml TGFβ (R&D Systems #240-B) + 10 ng/ml EGF.

#### Collagen coating protocol

Eight-well plates were coated with 20 μg/cm^2^ collagen-1 in a mixture that mimicked the buffering and structural characteristics of MEMA spots: 200 μg/ml collagen-1 (Cultrex #3442-050-01), 10% v/v glycerol (Sigma G5516), 5 mM EDTA pH 8 (Invitrogen 15575), and 100 mM Tris-HCl pH 7.2 (Sigma T2069) in PBS. Plates were rocked at RT for 1 hour.

Remaining coating mixture was gently aspirated and plates were washed twice with sterile PBS. Wells were allowed to dry completely by leaving the plate uncovered in a laminar flow hood before being stored in a benchtop desiccator for a minimum of three days and maximum of six months before use.

#### Data collection batches

Samples were collected over three collection periods. The first collection was completed at OHSU in the Fall of 2017 when RPPA, RNAseq, ATACseq, L1000, and IF samples were collected. The second collection was completed at OHSU in the Winter of 2018 and included GCP, L1000, and IF samples. The third collection was collected at HMS in the Summer of 2018 and included cycIF and L1000 samples.

### MCF10A Dose Optimization

MCF10A cells were plated on collagen coated 24-well plates in full growth media for 7 hours at which point the media was exchanged for experimental media. Following 18 hours in experimental media, fresh experimental media was added with 7 doses of OSM, EGF, and HGF individually, or with seven doses of BMP2, IFNG, and TGFB in combination with 10ng/ml EGF. After 72 hours in ligand containing media, cells were fixed, stained with DAPI, and imaged on the ScanR microscope. Cell counts from the images were quantified using Cell Profiler and normalized based on the number of cells present in the 10ng/ml EGF condition.

### OSM validation experiments

To assess responses to JAK/STAT inhibition MCF10A cells were plated in 24-well collagen coated plates. Following the media changes, cells were treated with 10 ng/ml OSM, 10 μM ruxolitinib (Selleck Chemicals #S1378) and Nuclight Rapid Red Dye (Essen Bioscience #4717) to label nuclei and count cells across time. Cells were then placed in the IncuCyte S3 and imaged every 30 minutes for 48 hours using phase contrast and red fluorescent filter sets. Cell number was quantified in Cell Profiler by counting the number of fluorescent nuclei in each frame and normalizing counts to time 0H.

To assess cell responses to protease inhibitors cells were plated in 24-well collagen coated plates, underwent the standard media changes and then at time 0H treated with 10 ng/ml OSM and either a protease inhibitor cocktail at a 1:400 dilution (Sigma-Aldrich #P1860), 40 μM bestatin (Sigma-Aldrich # B8385), 800 nM aprotinin (Sigma-Aldrich # A1153), 10 μM E-64 (Sigma-Aldrich # 324890), 1.45 μM pepstatin (Sigma-Aldrich # P5318). Cells were then placed in the IncuCyte S3 and imaged every 30 minutes for 48 hours.

Phase contrast images were registered using a custom ImageJ script and then imported into the Baxter Algorithms cell tracking software^89^. Clusters of cells with an area greater then 1000 pixels (~5 cells) were tracked using default parameters. Cell cluster tracks were then analyzed to quantify migration. Speed, displacement, mean squared displacement, and the cumulative distance traveled was calculated for cell clusters.

### Live-cell imaging

Well plates were placed in the IncuCyte FLR and phase contrast images were acquired every 30 minutes for 48 hours. Individual cells were manually tracked using the Fiji^90^ plugin MtrackJ^91^. Custom R scripts were used to quantify the migratory behavior of individual cell lineages. In brief, starting at the last time slot of each lineage, one cell was randomly selected and traced back through mitotic events until T0. Migration distance for each lineage was then calculated as the sum of the distances in pixels along the path between each image. To compare migratory behavior across different ligand treatments, we performed an ANOVA followed by Tukey’s Honestly Significant Difference test in R. Ligand treatments with p-value < 0.05 were deemed significantly different.

### Immunofluorescence

Prior to fixation, cells were pulsed with 10 μM EdU (Thermo Fisher Scientific C10357) for 1 hour under standard culture conditions. Cells were then fixed for 15 minutes with 2% paraformaldehyde (Electron Microscopy Sciences #15710) and permeabilized for 15 minutes with 0.01% Triton X-100 in PBS. Cells were then stained with CellMask (Thermo Fisher Scientific #H32713) for 30 minutes at RT, followed by fluorescent labeling of incorporated EdU for 1 hour at RT (Thermo Fisher Scientific C10357). Finally, cells were stained with a keratin 5 polyclonal antibody (BioLegend #905501) at 1:800 overnight at 4°C, followed by an anti-rabbit 488 secondary antibody (Thermo Fisher Scientific A21206) at 1:300 and Dapi (PromoKine PD-CA707-40043) at 0.5 μg/μL for 1 hour at RT.

Fixed cells were imaged on an Olympus ScanR microscope. DAPI channel images were imported into Ilastik for pixel classification^92^. A set of 20 images per plate were randomly selected and used for training. Pixels were classified as either nuclei or background using all default intensity, edge, and texture features, and with smoothing filters ranging from 0.3 – 10 pixels. Probability maps were then exported from Ilastik into CellProfiler version 3.1.8 for object segmentation^93^. Nuclei were identified using the global Otsu method with a threshold smoothing scale of 1.35. Clumped nuclei were separated based on intensity with a smoothing filter of 12 pixels. Cytoplasm compartments were assigned to nuclei by a 10-pixel donut expansion from each nucleus. Cytoplasm and nuclear Intensity, size, and morphology data was then exported into RStudio (RStudio Team, 2015). The values are analyzed as populations that have been median summarized from the cell-level data to the image or field level. The field level data are then median summarized to the well level. The EGF time course normalized values are the raw values divided by the corresponding EGF value at the same time point within the same replicate set. The preprocessing and QA script is at https://github.com/MEP-LINCS/MDD/tree/master. All samples passed qualitative QC inspection that the integrated DAPI intensity has the expected bimodal distribution.

### Phenotype analysis

All phenotypic quantifications were derived from immunofluorescent cell-level data. *Cell cycle phase* was determined by analysis DAPI intensity: each cell was classified into either G1 or G2M cell cycle phase by clustering cells into two groups based on total nuclear DAPI intensity. The Forgy k-means algorithm was used for clustering (R stats package), with the number of centers set to two. DAPI thresholds for classification were manually inspected, and multinucleated and poorly segmented cells were removed from further cell cycle analysis. *KRT5 intensity* was calculated as the mean intensity value of KRT5 in the cytoplasmic cell compartment.

Three spatial metrics were computed to quantify treatment induced changes in cell clustering and dispersal. The *number of neighbors* for each cell was calculated by quantifying the number of cell centroids within 100 pixels of a cell’s centroid. Cells with coordinates less than 100 pixels from the image border were excluded. *Nearest neighbor distances* were determined by measuring the pixel Euclidean distances of each cell centroid to the centroids of the four nearest cells in the imaging field. To account for variations in image cell count, the mean nearest neighbor distances for each image were normalized by the expected mean distance to the nearest neighboring cell if the cells were distributed randomly^94^. The *number of cells per cluster* was computed in a two-step process: first performing mean shift clustering on the cell centroid coordinates for each image, using the R package LPCM (v 0.47), and then computing the mean number of cells per cluster.

To compare phenotypic responses across treatments, we analyzed quantifications of the immunofluorescent images 48 hours after treatment. The Kruskal-Wallis test was used to test for overall treatment dependent differences. Pairwise comparisons between treatments were then conducted using Pairwise Wilcoxon Rank Sum Tests followed by an FDR p-value correction. For all tests, a q-value < .05 was considered significant.

### Reverse Phase Protein Array

#### Sample preparation

Cells were washed twice with ice-cold PBS followed by collection by manual scraping in 50-100 μL of lysis buffer (1% Triton X-100, 50mM HEPES pH 7.4, 150mM NaCL, 1.5mM MgCl_2_, 1mM EGTA, 100mM Na pyrophosphate, 1mM Na_3_VO_4_, 10% glycerol, 1x cOmplete EDTA-free protease inhibitor cocktail (Roche #11873580001), 1x PhosSTOP phosphatase inhibitor cocktail (Roche #4906837001)). Lysates were incubated on ice for 20 minutes with gentle agitation every 5 minutes and then centrifuged at 14,000 rpm for 10 minutes at 4°C. Supernatant was collected into a fresh tube, quantitated by BCA assay, and the appropriate volume was combined with 4X SDS sample buffer (40% glycerol, 8% SDS, 0.25M Tris-HCl, 10% ß-Me, pH 6.8), boiled for 5 minutes, and stored at −80°C. Three sets of replicates were collected over three weeks and submitted to MD Anderson Cancer Center for RPPA testing.

#### Pre-processing and QC

Samples underwent standard pre-processing using methods developed at the MD Anderson Cancer Center RPPA core^95^. In brief, the processing steps include the following: 1) Convert raw data from log2 value to linear value. 2) Determine median for each antibody across the sample set. 3) Calculate the median-centered ratio by dividing each raw linear value by the median for each antibody. 4) Assess sample quality by computing a correction factor (CF.1) for protein loading adjustment for each sample as the median of the median-centered ratio values from Step 3 for all antibodies. Samples with correction factors above 2.5 or below 0.25 are considered outliers and discarded. 5) Compute the normalized linear value by dividing the median-centered ratio from Step 3 by CF.1. All samples passed MDACC’s quality checks and are included in the dataset. The normalized RPPA log2 values are joined with their experimental metadata and stored on Synapse as level 3 data. Replicates are median summarized and stored as Level 4 data.

### RNA Sequencing

#### Sample preparation and sequencing

Following treatment protocols described, at the appropriate timepoint wells were aspirated and cells were harvested by scraping in 600 μl of

RLT Plus buffer (Qiagen) plus 1% β-ME. Samples were flash frozen in liquid nitrogen and stored at −80°C prior to RNA extraction. Total RNA was extracted from frozen using a Qiagen RNeasy Mini kit. Columns were DNAse treated following the recommended protocol of the manufacturer.

RNA concentration and purity was determined by UV absorption using a Nanodrop 1000 spectrophotometer. All samples had 260/280 absorption ratios of at least 2.0, indicating successful isolation of RNA from other nucleic acids. RNA integrity was assessed using an Agilent 2100 Bioanalyzer with an RNA 6000 Nano Chip. RNA integrity numbers (RIN) were calculated from Bioanalyzer electropherograms using the “Eukaryotic Total RNA Nano” program of the Bioanalyzer 2100 Expert software (B.02.08.SI648). RIN values were in the 8.5-10 range, indicating high-quality RNA, with one exception (TGFB_48_C1_B; RIN = 6.9). UV absorption measurements and RIN values are available on Synapse (https://www.synapse.org/#!Synapse:syn12550434).

cDNA libraries were prepared from polyA-selected RNA using an Illumina TruSeq Stranded mRNA library preparation kit. 100-bp single-end reads were sequenced on an Illumina HiSeq 2500 Sequencer, with a target of 60M reads per sample.

#### Pre-processing and QC

Sequence preprocessing and alignment was performed using a Docker-based pipeline^96^. 100-bp single-end reads were trimmed of Illumina adapter sequences using TrimGalore (v. 0.4.3), a wrapper for CutAdapt (v. 1.10) and FastQC (v. 0.11.5). A minimum of 1-bp overlap with the adapter sequence (AGATCGGAAGAGC) was required for trimming. After trimming, reads with a length < 35 bp were discarded. Trimmed reads were aligned to the GENCODE V24 (GRCh38.p5) assembly of the human genome using the Kallisto pseudo-alignment software (v. 0.43.0). Kallisto, using the following parameters: --bias-b 30 --pseudobam.

Gene-level quantifications were produced from transcript-level abundance estimates using the R (v. 3.5.0) package tximport (v. 1.8.0). Mapping between gene/transcript identifiers was done using the biomaRt package (biomaRt v. 2.36.1) with the ENSEMBL_MART_ENSEMBL biomart and the hsapiens_gene_ensembl dataset. Gene-level quantifications were imported to DESeq2 (v. 1.24.0)^97^. The fpkm function of DESeq2 was used to normalize data for library size and gene length differences, and fpkm values were log2 transformed with an added pseudocount of 1.

#### Transcription Factor Enrichment Scores

Single-sample enrichment scores were calculated for 297 transcription factor target gene sets obtained from the CHEA3 ReMap_ChIP-seq^68^ using the R package GSVA (v. 1.32.0)^98^. A minimum expression filter was used to filter for expressed genes; genes were retained only if expressed at a minimum of 0.5 log2(fpkm + 1) in a minimum of 3 samples. Enrichment scores were calculated from filtered RNAseq data, in units of log2(fpkm + 1), using the argument “method = ‘ssGSEA’”.

### ATACseq

#### Sample preparation and sequencing

ATACseq samples were collected following the Omni-ATAC protocol^99^. Briefly, MCF10A cells were washed once with PBS and detached from the plate using trypsin. Cells were then counted using a Countess (Invitrogen), and 50,000 cells per condition were distributed to 1.5 ml centrifuge tubes and spun at 500 RCF for 5 min. The supernatant was removed and the cell pellet was resuspended in 500 μl of PBS and spun again at 500 RCF for 5 min. The supernatant was removed again, and the cell pellet was resuspended in 50 μl of cold ATAC resuspension buffer (RSB) containing 0.1% NP40, 0.1% Tween-20, and 0.01% digitonin by pipetting up and down three times. After 3 min on ice, 1 ml of cold RSB containing 0.1% Tween-20 was added, and the tube was inverted three times to mix. The nuclei were then pelleted by centrifugation at 500 RCF for 10 min at 4°C. The supernatant was then carefully aspirated and the nuclei were resuspended in 50 μl of transposition buffer (25 μl 2x TD buffer (Illumina), 2.5 μl transposase (Illumina), 16.5 μl PBS, 0.5 μl 1% digitonin, 0.5 μl 10% Tween-20, and 5 μl H2O). Samples were then placed in a pre-warmed (37°C) thermomixer and mixed for 30 min at 100 RPM. Transposed fragments were then purified using a Qiagen MinElute column and frozen at −80°C for further processing.

The remaining steps of the Omni-ATAC protocol were performed by the OHSU Massively Parallel Sequencing Shared Resource. Transposed fragments were pre-amplified with 5 rounds of PCR. Afterward, 5 μl of the pre-amplified mixture was used for a qPCR reaction to determine the concentration of tagmented DNA. After calculating the concentration of tagmented DNA, pre-amplified samples were diluted with elution buffer to a final concentration of 5 μM. Six samples had an undiluted DNA concentration below 5 μM and were not diluted. 5 μM preamplified samples were amplified for 3 additional PCR cycles.

Tagmented DNA was pre-amplified with 5 rounds of PCR (72°C for 5 min, 98°C for 30 seconds, then 5 cycles of [98°C for 10 sec, 63°C for 30 sec, 72°C for 1 min]). PCR reactions contained 20 μl eluate, 25 μl NEBNext 2x MasterMix, 2.5 μl 25 μM i5 primer and 2.5 μl 25 μM i7 primer.

The DNA concentration of the pre-amplified samples was assessed by qPCR. 5 μl of preamplified mix was added to 3.76 μl sterile water, 0.5 μl 25 μM i5 primer, 0.5 μl 25 μM i7 primer, 5 μl 2x NEBNext master mix, and 0.24 μl 25x SYBR Gold (in DMSO). Samples were amplified for 20 cycles of [98°C for 10 sec, 63°C for 30 sec, 72°C for 1 min]. DNA concentration was calculated, and pre-amplified samples were diluted to a final concentration of 5 μM. Six samples had an undiluted DNA concentration below 5 μM and were not diluted. 5 μM pre-amplified samples were amplified for 3 additional PCR cycles. 100bp PE reads were sequenced on an Illumina HiSeq 2500 Sequencer by the OHSU Massively Parallel Sequencing Shared Resource with a target of 20M reads per sample.

#### Pre-processing and QC

ATACseq files were processed and aligned using the “ATACseq (1 -> 3)” workflow on the AnswerALS Galaxy server (answer.csbi.mit.edu). Reads were trimmed of adapter sequences and low-quality bases using Trimmomatic (Galaxy version 0.36.5). Reads were trimmed of low-quality bases (Phred score < 15) at the read start or end, and Nextera adapter sequences (CTGTCTCTTATA) were trimmed from read ends (minimum of a 2-bp overlap required for trimming). Reads were aligned to the human genome (hg38) using Bowtie2 (Galaxy version 2.3.4.1) in paired-end mode with otherwise default settings. BAM files were filtered to remove secondary alignments, unmapped reads, and mitochondrial DNA alignments using ngsutils bam filter (Galaxy version 0.5.9). PCR duplicates were detected and removed using Picard MarkDuplicates (Galaxy version 2.7.1.2). The de-duplicated, filtered BAM file was used for peak calling and quantification. Peaks were called using MACS2 (Galaxy Version 2.1.1.20160309.5) using the following parameters: -format BAMPE -nomodel -extsize 200 -shift −100 -qvalue 0.01.

ATACseq sample quality was assessed by calculating the fraction of reads in peaks (FRiP). Before calculating FRiP, a consensus peakset was generated for all samples by taking the union of all peaks called in all samples and merging any overlapping peaks, using the R (v. 3.6.1) package DiffBind (v. 2.12.0)^100^. For each sample, FRiP was then calculated by counting the proportion of reads in the de-duplicated, filtered BAM file that align within the consensus peakset. A minimum FRiP threshold of 0.15 was applied to remove samples with low levels of chromatin enrichment.

#### Construction of chromatin accessibility matrix

DiffBind (v. 2.12.0) was used to generate a peak accessibility matrix for the QC-passing samples. First, a consensus peakset was re-generated after removal of low-FRiP samples. The dba.count function was then used to count the number of reads in the de-duplicated, filtered BAM files that overlap with each peak in the consensus peakset. The dba.count argument “score = DBA_SCORE_TMM_READS_EFFECTIVE” was used to output TMM counts normalized to each sample’s effective library size, which is equal to the de-duplicated, filtered library size multiplied by the FRiP. A peak accessibility matrix in units of unnormalized counts was also generated using the dba.count function with the argument “score = DBA_SCORE_READS”.

#### Motif Enrichment

Transcription factor motif enrichment scores were generated from the TMM-normalized chromatin accessibility data using the R package chromVAR (v. 1.6.0)^101^. ATACseq peaks were annotated with GC content using the addGCBias function of chromVAR and the BSgenome.Hsapiens.UCSC.hg38 genome annotation package. Transcription factor motif position frequency matrices were obtained from the “JASPAR CORE 2018 Homo sapiens” set of motifs^102^. ATACseq peaks were matched to JASPAR motifs using the R package motifmatchr (v. 1.6.0). The expected fraction of reads per ATAC-seq peak was calculated using the chromVAR function computeExpectations, with the argument “norm = TRUE”. Each sample’s deviation from the expected fraction of peaks in each annotated category was calculated using the function computeDeviations, and deviations were converted to Z-scores using the function deviationScores.

### Global Chromatin Profiling

The GCP assay was performed as previously described in Creech *et al*^51^ and Litichievskiy *et al*^11^ Cells were washed with ice-cold PBS, then collected by manual scraping in 200 μl of cold PBS. Cells were then pelleted by centrifugation at 1500 RCF at 4°C for 5 min, resuspended in 1mL of cold PBS, and spun again as specified. The resultant cell pellets were then flash frozen in liquid nitrogen and stored at −80°C until further processing. Pellets were thawed and lysed with nucleus buffer, followed by histone extraction by sulfuric acid and precipitation using trichloroacetic acid. Sample input was normalized to 10 μg of histone in H_2_O before being propionylated, desalted (Oasis HLB 5mg Plate) and digested by Promega trypsin overnight. A second round of propionylation, followed by desalting using C18 Sep-Pak cartridges (Waters) was employed after digestion. Propionylations and digestion were done in an automated fashion on an LT-Bravos system (Agilent). Isotopically labeled synthetic peptides from histones H3 and H4 were added as a reference to each sample prior to MS analysis. Peptides were separated on a C18 column (EASY-nLC 1000, Thermo Scientific) and analyzed by MS in a PRM mode (Q Exactive™-plus, Thermo Scientific) as previously described^51^. Detailed protocols of sample preparation steps can be found in https://panoramaweb.org/labkey/wiki/LINCS/Overview%20Information/page.view?name=sops.

### L1000

#### Sample preparation

L1000 samples were collected as part of three collections. The first L1000 sample collection was generated in parallel to the ATACseq samples. MCF10A cells were washed once with PBS and detached from the plate using trypsin. Cells were then counted using a Countess (Invitrogen) and 50,000 cells per condition were distributed to 1.5 ml centrifuge tubes and spun at 500 RCF for 5 minutes. The supernatant was removed and the cell pellet was resuspended in TCL buffer (Qiagen) containing 1% β-Me. For the second and third collections, cells were washed with PBS followed by the addition of TCL buffer (Qiagen) containing 1% β-Me. The cell and buffer mixture was allowed to sit for 30 minutes and then frozen at −80°C for further processing. Samples from the first and second sample collections were frozen in 1.5ml tubes. Samples from the third data collection were frozen in their original 96-well plates. In total there were eighteen plates from the third HMS collection, which contained 21 samples per plate, and there were 190 samples from the first two OHSU collections. All samples were shipped to the BROAD for simultaneous processing on the L1000 platform. The source plates containing original samples were re-arrayed into six 96-well master plates. These master plates contained 21 samples from each of three original source plates, and 32 samples plated directly from tubes. In each of the six master plates, well A1 was left empty to accommodate for internal technical control spike-ins. The six 96-well master plates were then re-arrayed into the final 384 well v-bottom PCR Plates (Eppendorf #951020702).

#### Ligation Mediated Amplification

Complete methods for L1000 Ligation Mediated Amplification can be found elsewhere^20^. In brief: crude cell lysates were transferred from source plates to 384 well v-bottom PCR Plates (Eppendorf #951020702) assay plates. Oligo dT coated magnetic particles (GE Healthcare #38152103010150) were added to capture mRNA. Plates were then incubated at room temperature on shaker tables for 10 minutes. The beads were then spun down onto flat magnets and unbound lysate was evacuated by centrifuging upside down on magnet to 800RPM for 30 seconds. 15μl of reverse transcription master mix containing SuperScript IV reverse transcriptase was added to the plates and the plates were incubated at 55 °C for 10 minutes. Plates were again spun down, beads were pelleted on a flat magnet, and the remaining master mix was spun out. Probes were annealed to the first-strand cDNA by addition of 15μl of Probe Bind master mix, containing 100 fmole of each probe and Taq ligase buffer. Samples were denatured at 95 °C for 5 minutes, then transferred to a ramping water bath that decreased temperature from 70 °C to 40 °C over six hours. The following day, beads were again spun down on a flat magnet and master mix was evacuated. To ligate probe pairs, 15 μL of Ligation Master Mix was added, containing Taq DNA ligase and ligase buffer. Plates were sealed and incubated at 45°C for 60 minutes. Plates were spun down on magnets and ligation master mix was evacuated as with previous steps. 15μl PCR master mix containing 0.5 mmole of each primer (T3 and 50-biotinylated T7 universal primers), dNTPs, and PlatinumTaq polymerase in reaction buffer was added to each well, and plates were subjected to 29 cycle PCR. This process yielded biotinylated gene and bead (barcode) specific amplicons.

Each barcode corresponds to a complementary sequence on a Luminex bead, allowing the PCR product to be hybridized to a mixture containing per well ~100 each of 500 Luminex analyte colors. The plate was then denatured at 95°C for 5 minutes and incubated at 45°C for 18 hours. Beads were pelleted and stained with streptavidin R-phycoerythrin conjugate for ten minutes. Finally, plates were read on Luminex FlexMap 3D Flow cytometers that detected analyte color (transcript identity) and fluorescence intensity (transcript abundance) for all analytes detected in all wells.

#### Pre-processing

To account for differences across the various cell collections, we adapted our standard data processing pipeline in several ways. L1000 data typically use a population-based normalization scheme, known as plate control, as described in Subramanian *et al*^20^. Here, the EGF treated wells served as the vehicle when conducting vehicle normalization. The standard data processing pipeline was followed, except for the changes at Level 1 and Level 4, described below.

L1000 utilizes 10 sets of invariant genes, similar to ‘housekeeping’ genes, to assess quality and in later normalization steps. These gene sets, each containing 8 genes, represent control values that span the spectrum of gene expression, and are ordered according to their overall level of expression, the first level corresponding to the lowest expressing genes, and the 10th corresponding to the highest expressors.

Plates were computationally split at Level 1 (LXB) into subpopulations of wells, each containing only samples from a given time-point and collection combination. The fluorescence intensity values associated with each bead color were subjected to the peak deconvolution step, which separates the two genes associated with each bead color (Level 2). Data were then normalized via L1000 invariant set scaling (LISS), which scales the expression levels of the 978 measured landmarks in each well to the 80 control genes in the invariant gene set (Level 3). Next, we calculated differential expression using EGF as the vehicle control. Robust z-scoring was used to calculate differential expression values for each gene, where gene x is compared only to the vector of normalized gene expression of gene x across all EGF samples in that collection/time-point population (Level 4). Finally, individual biological and technical replicates were collapsed into a consensus signature by computing a pairwise Spearman correlation matrix between each replicate signature. The weights for each replicate were calculated by the sum of their correlations to the remaining replicates, summing to 1. The consensus signatures were generated by the linear combination of the replicate signatures using each signature’s weight as the coefficient (Level 5).

#### L1000 QC

We used several approaches to assess data quality. First, to assess the quality in each detection plate, we visually inspected and measured the slope of the invariant gene calibration curve for each sample; outliers were omitted. Second, to assess plate effects, we plotted median fluorescence intensity and interquartile range of invariant set 10 across the entire plate. This allowed identification of failed (low signal) wells, tissue culture related plate effects, or wells with abnormally wide ranges in expression across each gene set. Third, to assess the efficacy of the deconvolution algorithm, we determined the number of well/analyte combinations where two peaks were clearly discernible.

In addition, we computed a transcriptional activity score (TAS) as a composite measure of L1000 transcriptional response. Here signature strength (SS) was computed as the number of genes with a z-score greater than or equal to 2 for each sample, and replicate correlation (CC) was computed as the 7th quantile of the spearman correlation between all pairwise combinations of replicates. TAS is calculated as the geometric mean of SS and CC for a signature, and scaled by the square root of the number of landmark genes, yielding a final score between 1 and 0. QC metrics are available on Synapse (https://www.synapse.org/#!Synapse:syn19416843).

Finally, within each sample collection (C1, C2, and C3), we clustered samples based on the Euclidian distances between expression of the 978 measured landmark genes in the Level 3 data, using the R function hclust. Each collection had a small number of outlier samples that showed markedly aberrant expression of the 978 landmark genes and clustered apart from all other samples, in a pattern that was not explained by sample treatment; these samples were removed. In total, 17 L1000 samples were removed (3 from C1, 1 from C2, and 13 from C3).

### Cyclic Immunofluorescence (CyCIF)

#### Sample preparation and imaging

MCF10A cells were seeded 4000 cells/well in 200 μl of GM in collagen coated (as described above) 96 well plates (NUNC, 165305) in technical (multiple wells on the same plate) and biological (experiments separated by a minimum of one cell passage) triplicates. Eight hours after seeding, the cells were washed once with PBS using an EL405x plate washer (BioTek), and 200 μl of EM was added per well. Following an additional 16 hours (24 hours after initial plating), one plate was fixed (time = 0 hours) and EM was aspirated from all wells in the remaining plates using the plate washer, and replaced with 200 μl of the appropriate ligand or control treatment.

The treated plates were fixed following incubations of 1, 4, 8, 24, and 48 hours. Cells were fixed in 4% formaldehyde for one hour at room temperature, and washed with PBS. Plates were sealed and stored at 4°C until all replicates were collected. CycIF was performed as described previously^47,48^. In brief, cells were permeabilized with ice cold methanol for ten minutes, blocked in Odyssey buffer (LI-COR) for one hour, pre-stained with secondary antibodies, bleached, and imaged to register background intensities prior to beginning cycIF. For each cycle, cells were stained with three conjugated antibodies, unless otherwise specified, and Hoechst 33342 overnight at 4°C, washed with PBS, and imaged with an IN Cell Analyzer 6000 (nine fields of view per well, 20x/0.45NA air objective, 2൸2 binning) (GE Healthcare Life Sciences). Following image acquisition, fluorophores were chemically inactivated as described^47,48^, and cells then entered the next staining cycle. Refer to **Supp Table 12** for antibody metadata.

#### Pre-processing and image analysis

A flat field correction profile, generated from all fields on one plate using the BaSiC ImageJ plugin^103^, was normalized to a mean value of one and each image was then divided by it. Image registration was performed with a custom ImageJ script. Segmentation of the nuclei (based on Hoechst staining), and cytoplasm (based on β-catenin staining) was performed with a custom MATLAB (MathWorks) script. Each cell was then divided into four subcellular masks: nucleus, peri-nuclear ring, cytoplasm, and cell membrane for feature extraction, a fifth region including all of the cytoplasm (peri-nuclear ring, cytoplasm, and cell membrane together) was also defined. Segmentation was performed on the images acquired in cycle 4 only; the masks were then overlaid on all other cycles for feature extraction. Intensity, texture, and morphology features were extracted for each mask, as appropriate (see **Supp Table 13** for feature definitions).

#### CycIF QC

Quality control was performed in two steps. In the first step, cells that were washed away over the course of the experiment and those near the edges of the imaging fields that were incompletely captured cycle to cycle due to microscope stage drift were identified and excluded from subsequent analyses. These cells were identified by their high variation in nuclear Hoechst signal between successive cycles (https://github.com/yunguan-wang/cycif_analysis_suite/blob/MCF10A/notebooks/Section2.1_Intensity%20based%20QC.ipyn_b). If more than 90% of the cells in a field of view failed this QC step, the entire field was removed. The median fraction of lost cells was ~15 % for fields 1-8 whereas an average of 60% of cells were lost from field 9, with a significant number of instances where the fraction of lost cells exceeded 90%. Field 9 was therefore excluded entirely from subsequent analyses. Additionally, for unknown reasons, most of the wells occupying row E on plate 18 exhibited cell loss in excess of 90% leading to the exclusion of all data from those wells in downstream analyses. In the second quality control step, cells with failed cytoplasm segmentation as identified by multinucleation were removed. Multi-nucleated cells were identified by resegmenting each mask using the Python implementation of Opencv (https://github.com/skvark/opencv-python) and counting the nuclei; cells with two or more nuclei were excluded from downstream analyses (https://github.com/yunguan-wang/cycif_analysis_suite/blob/MCF10A/notebooks/Section2.2_image_based_qc.ipynb). Although masks with two nuclei can represent failed segmentation or truly binucleated cells, visual inspection led us to conclude that these cases were primarily segmentation errors and were therefore excluded from downstream analyses.

### Identification of differentially-expressed genes

For each ligand treatment, we performed a differential expression analysis on the RNAseq gene-level summaries with the R package DESeq2 (1.24.0), with shrunken log2 fold change estimates calculated using the apeglm method. We used the Benjamini-Hochberg method to correct p-values for multiple comparisons and set a threshold of q-value < 0.05 and shrunken log2 fold change > 1.5 or < −1.5 to indicate significance.

### Pathway enrichment of ligand-induced signatures

We used Gene Set Enrichment Analysis (GSEA) to identify the pathways enriched by each ligand treatment. Specifically, we used Gene Set Enrichment Analysis 4.1.0 downloaded from https://www.gsea-msigdb.org/gsea/index.jsp to assess enrichment of the MsigDB Hallmark Pathways in the Level 3 data. For each 24H ligand treatment sample, we computed log2 fold-change against CTRL_0 from the Level 3 RNAseq data.

### L1000 drug signature comparison

To compare our results to existing L1000 transcriptional drug signatures^20^ we used the L1000 FWD tool^104^ available at https://maayanlab.cloud/L1000FWD/. We used as input the top 200 most strongly up-regulated and top 200 most strongly down-regulated genes at 24 H relative to CTRL_0. We considered drug signatures with Fisher exact test q-values < 0.2 to be significantly correlated or anticorrelated with our ligand signatures. Finally, we summarized the number of drugs with similar mechanisms of action to identify common patterns.

### Multi-omic module detection

To identify coordinately regulated multi-omic modules, we performed normalization, data scaling, feature selection and cluster analysis on molecular features induced by ligand treatments.

#### Data normalization and scaling

For the GCP, RPPA and cycIF datasets we used limma to normalize to CTRL_0 and summarize across the replicates; we used DESeq2 to analyze the RNAseq data in a similar manner. We used chromVAR to aggregate chromatin accessibility peaks that share common motifs and then mean summarized the motif family values. We applied the rrscale transformation to each assay data set to minimize assay-specific data distributions^67^. In brief, each assay’s T0 CTRL normalized data was rrscaled independently with Box Cox negative and asinh transformations using an infinite z score cutoff.

#### Feature selection

We selected a subset of highly variant and biologically interpretable features from the 24H and 48H samples from each assay. In GCP and RPPA assays, features in the lowest variance quartile were removed. For the cycIF, RNAseq, and GCP assays, features were retained if, for any condition, the absolute log fold change was greater than 1.5 and the p-value was less than 0.05. For the RPPA assay, we used a log fold-change threshold of 0.75 to account for differences in the RPPA data distribution. All ATACseq motif family scores were retained.

#### Clustering

We performed k-means clustering using partitioning around medoids and a gap statistic analysis using the firstSEmax method to identify the optimal number of clusters (R package cluster, version 2.1.2). In brief, the gap statistic method runs PAM clustering on the integrated data matrix once for each k value, where k=2:25. Then for each k, we performed PAM clustering on 100 randomized permutations of the data that have structure similar to the actual data. At each k, the gap is calculated as the difference in the log of the within-groups sum of squares of the actual versus randomized data. We further refined these clusters by identifying and collapsing highly correlated clusters. In brief, we calculated the mean expression of features in each cluster for each condition and then computed Pearson correlations between all pairs of clusters. Next, we then used the R hclust function and the dendextend cutree function on the distance matrix of the correlations to identify highly correlated clusters. This resulted in combining 4 pairs of clusters to yield a final set of 14 modules for further analysis.

### Module TF enrichment analysis

We identified transcription factors enriched in the integrated modules by submitting all RNAseq features from each integrated module to the ChEA3 webbased transcription factor enrichment tool ChEA3^68^, which identifies transcription factors enriched for a list of genes using Fisher’s exact test. We limited our analyses to transcription factor targets in the ReMap ChIP-Seq library and considered transcription factors significantly enriched if the FDR-corrected q-value was less than 0.2.

### Module pathway enrichment analysis

To identify pathways enriched in each module, we used the Reactome pathway enrichment analysis tool (https://reactome.org/) to analyze the genes in each module. In brief, this analysis performs a binomial test of each gene set of 2516 curated pathways in the Reactome database. We identified significantly enriched pathways as those with FDR q-values < 0.1, gene ratios > 0.1, and pathways that included a minimum of 5 and maximum of 500 genes.

### Module expression scores

To calculate the expression of modules across different samples in our MCF10A dataset, we computed the mean expression of features in each module. To assess expression of the modules in external datasets (*e.g*. GTEx), we focused on the RNAseq features in each module and computed their mean expression. An unpaired two-sample t-test was used to compare mean module 9 expression scores between control and CDKi-treated breast cancer cell lines.

### Set analysis

Set analysis was used to identify features significantly induced by a single ligand (ligand-specific) or multiple ligands (shared). The input to the set analysis was the integrated and scaled matrix of log fold change values derived from the multi-omic module analysis. Each feature in the multi-omic matrix was labelled either ‘Unique’ or ‘Shared’. Features that were significantly perturbed by only a single ligand were labelled ‘Unique.’ Features that were regulated by two or more ligands were labelled ‘Shared.’

## SOFTWARE

Unless otherwise stated, analyses were performed in R (https://www.R-project.org). R packages used in analyses included: tidyverse^105^ (version 1.3.1), ComplexHeatmap (version2.8.0), httr (version 1.4.2) and rmarkdown (version 2.9). A complete list of packages and their versions can be found in analysis scripts available at https://github.com/MEP-LINCS/MDD

## DATA AVAILABILITY

Data, metadata and additional analysis reports are available at: https://www.synapse.org/#!Synapse:syn21577710/wiki/601042. Raw RNA and ATAC sequencing data generated for this study can be accessed from the Gene Expression Omnibus (https://www.ncbi.nlm.nih.gov/geo/query/acc.cgi?acc=GSE152410).

## ACKNOWLEDGEMENTS

This work was funded by grants U54-HG008100 to JWG, LMH, and JEK; U54HL127365 to PKS; U54HG008098 to MRB; U54NS091046 to EF; R01-GM104184 to MRB; U54HL127624 to SS and AM; U54-HG008097 to JDJ; U54HL127366 to AS. CE was a LINCS Consortium Postdoctoral Fellow. NCI Cancer Center Support Grant P50CA16672 supported generation of RPPA data. OHSU GPSR and MPSSR receive support from the OHSU Knight Cancer Institute NCI Cancer Center Support Grant P30CA069533.

## AUTHOR CONTRIBUTIONS

**Conceptualization:** Laura Heiser, Joe Gray, Ajay Pillai, Albert Lee

**Study coordination and supervision:** Laura Heiser

**Cell culture:** Sean Gross, Kaylyn Devlin, Rebecca Smith, Tiera Liby, Moqing Liu, Joe Gray, Laura Heiser, James Korkola

**Immunofluorescence:** Sean Gross, Kaylyn Devlin, Rebecca Smith, Ian McLean, Mark Dane, Laura Heiser

**Live-cell imaging:** Sean Gross, Ian McLean, Crystal Sanchez-Aguila, Laura Heiser

**CycIF:** Caitlin Mills, Kartik Subramanian, Yunguan Wang, Connor Jacobson, Clarence Yapp, Mirra Chung, Peter Sorger

**MEMA:** Kaylyn Devlin, Rebecca Smith, David Kilburn, Mark Dane, James Korkola

**RPPA:** Yiling Lu, Mark Dane, Gordon Mills

**RNAseq:** Sean Gross, Daniel Derrick, Denis Torre, Avi Ma’ayan, Joe Gray, Laura Heiser **ATACseq:** Sean Gross, Daniel Derrick, Jonathan Li, Miriam Adam, Brook Wassie, Laura Heiser, Ernest Fraenkel

**L1000:** Nicholas Lyons, Ted Natoli, Sarah Pessa, Xiaodong Lu, Aravind Subramanian

**GCP:** James Mullahoo, Malvina Papanastasiou, Jake Jaffe

**Integrative analyses:** Mark Dane, Sean Gross, Daniel Derrick, Cemal Erdem, Alexandra London, Marc Birtwistle, Avi Ma’ayan, Joe Gray, Laura Heiser

**Data curation:** Mark Dane, Daniel Derrick, Elmar Bucher, Kenneth Daily, Dusica Vidovic, Larsson Omberg, Stephan Schurer

**Project Management:** Heidi Feiler

**Writing:** Sean Gross, Mark Dane, Ian McLean, Joe Gray, Laura Heiser. All authors reviewed and edited the manuscript.

---

**Figure S1. Experimental and bioinformatic approaches to identify high impact ligands**

(A) Microenvironmental assay (MEMA) to identify ligands that modulate MCF10A cell numbers. Cells were treated with ligands in experimental media lacking EGF and cell numbers were counted after 72H. (B) MEMA assay results for MCF10A cells treated with ligands in experimental media containing EGF. (C) MCF10A transcript expression from three receptor classes: Tyrosine kinase, cytokine, and TGFB/BMP. Transcript values are drawn from RNAseq measures for untreated cells in exponential growth. The primary receptors for the six ligands are highlighted HGF: MET (Blue), EGF:EGFR/ERBB2 (Red), BMP2:BMPR1B/BMPR1A (Green), TGFB: TGFBR1/TGFBR2 (Yellow), OSM: IL6ST/OSMR (Orange), and IFNG:IFNGR1/IFNGR2 (Purple). (D) Cell count dose-responses after treatment with EGF, HGF, and OSM. Cell counts at 72H were normalized to the 10 ng/ml EGF condition. (E) Cell count dose responses for TGFB1, IFNG, and BMP2. Each of the ligands were supplemented with 10 ng/ml EGF. Cells counts at 72H were normalized to the EGF condition with no secondary ligand.

**Figure S2. Comparison of ligand and small molecule inhibitor signatures**

We leveraged the LINCS L1000 database^57^ of drug response signatures to identify targeted inhibitors that are shared by each ligand signature. Heatmap represents the number of compounds that have correlated (red) or anti-correlated (blue) signatures with each ligand (Fisher exact test, q-value<0.2). The ligand panel activated many of the same signatures as small molecule inhibitors, indicating that shared molecular responses can be elicited by these distinct perturbagen classes.

**Figure S3. IFNG responses are dynamically encoded across multiple molecular modalities**

(A) Cartoon of canonical STAT pathway activation after treatment with IFNG ligand. (B) Line graphs show induction of pSTAT1, IRF1 and PDL1 protein expression following IFNG treatment, as measured by cycIF and RPPA assays. (C,D) Cyclic immunofluorescence images show changes in STAT1 and PDL1 protein abundance and localization induced by IFNG+EGF treatment. (E) Line graphs show enrichment of STAT-family and IRF-family motifs inferred from ATACseq chromatin accessibility data for IFNG+EGF and EGF conditions. (F) Chromatin accessibility near the IRF1 and PDL1 gene loci. The local gene region for IRF1 showed a new peak in the promoter region and a large accessibility change in the 3’ region. IFNG did not induce new ATAC peaks in PDL1, however IFNG induced a new peak in the adjacent PDL2 gene (PDCD1LG2). DNA regions with changes in accessibility are marked with a red background. (G) Venn diagram showing the overlap between the Reactome IFNG pathway, curated cytokine gene lists and, and genes induced by IFNG+EGF treatment. The 15 cytokines induced by IFNG+EGF are listed on the right.

**Figure S4. Comparison of RPPA, RNAseq and ATACseq assays reveals concordance across molecular modalities in response to ligand treatment**

(A) Scatter plots of paired RPPA and RNAseq measurements, showing three classes of observed relationships: linear, ligand-specific, and no change. (B) Heatmaps show genes and proteins with significantly up- or down-regulated expression after ligand treatment. (Left): Heatmap of significantly up- or down-regulated genes assessed by RNAseq (FDR p ≤ 0.01; log2FC ≥ |1|). (Right): Heatmap of significantly up- or down-regulated proteins assessed by RPPA (FDR p < 0.01; log2FC ≥ |0.5|). (C) Euler diagram showing intersections of differentially expressed RPPA proteins and RNAseq genes. The majority of features that were induced in both assays showed concordant responses, defined as both modalities induced in the same direction. (D,E) Dot plot showing the relationship between ATACseq transcriptional start site (TSS) accessibility and gene expression in the CTRL and EGF 48H samples. Note the switchlike relationship between gene expression and accessibility at the TSS, as has been described previously. The horizontal dotted line indicates the threshold for a gene being defined as expressed.

**FigureS5. Integrated analysis methods**

Pre-processed data from each assay are summarized, filtered, and scaled before being combined into a single matrix. PAM clustering with gap analysis was used to identify the optimal number of clusters represented in the integrated data matrix, which resulted in 18 modules. Pearson correlation analysis was used to identify pairs of clusters that showed similar expression patterns; two pairs of modules were combined to yield a final set of 14 modules.

**Figure S6. Identification and characterization of integrative molecular modules**

(A) Bar plot showing the number of features for each assay included in the integrative modules; note log 10 scale. (B) Gap analysis used to identify the optimal number of modules. (C) Module correlation matrix showing Pearson correlation values. Highly correlated cluster pairs 8+17, 3+4, 6+16, and 10+15 were combined to yield 14 clusters. (D) Bar plot showing the distribution of features for each assay across modules.(E) Bar plot showing the mean module expression for each of the ligand treatments.

**Figure S7. GTEX Module expression analysis**

(A) Heatmap showing GTEX tissue expression of the 14 integrative molecular modules reveals tissue-specific expression, suggesting molecular programs that may be particularly important for mediating normal and diseased functions across tissues.

## REFERENCES

1 Heldin, C. H., Lu, B., Evans, R. & Gutkind, J. S. Signals and Receptors. Cold Spring Harb Perspect Biol 8, a005900, doi:10.1101/cshperspect.a005900 (2016).

2 Duronio, R. J. & Xiong, Y. Signaling pathways that control cell proliferation. Cold Spring Harb Perspect Biol 5, a008904, doi:10.1101/cshperspect.a008904 (2013).

3 Ward, P. S. & Thompson, C. B. Signaling in control of cell growth and metabolism. Cold Spring Harb Perspect Biol 4, a006783, doi:10.1101/cshperspect.a006783 (2012).

4 Devreotes, P. & Horwitz, A. R. Signaling networks that regulate cell migration. Cold Spring Harb Perspect Biol 7, a005959, doi:10.1101/cshperspect.a005959 (2015).

5 Perrimon, N., Pitsouli, C. & Shilo, B. Z. Signaling mechanisms controlling cell fate and embryonic patterning. Cold Spring Harb Perspect Biol 4, a005975, doi:10.1101/cshperspect.a005975 (2012).

6 Barretina, J. et al. The Cancer Cell Line Encyclopedia enables predictive modelling of anticancer drug sensitivity. Nature 483, 603–607, doi:10.1038/nature11003 (2012).

7 Costello, J. C. et al. A community effort to assess and improve drug sensitivity prediction algorithms. Nat Biotechnol 32, 1202–1212, doi:10.1038/nbt.2877 (2014).

8 Garnett, M. J. et al. Systematic identification of genomic markers of drug sensitivity in cancer cells. Nature 483, 570–575, doi:10.1038/nature11005 (2012).

9 Ghandi, M. et al. Next-generation characterization of the Cancer Cell Line Encyclopedia. Nature 569, 503–508, doi:10.1038/s41586-019-1186-3 (2019).

10 Heiser, L. M. et al. Subtype and pathway specific responses to anticancer compounds in breast cancer. Proc Natl Acad Sci U S A 109, 2724–2729, doi:10.1073/pnas.1018854108 (2012).

11 Litichevskiy, L. et al. A Library of Phosphoproteomic and Chromatin Signatures for Characterizing Cellular Responses to Drug Perturbations. Cell Syst 6, 424–443 e427, doi:10.1016/j.cels.2018.03.012 (2018).

12 Neve, R. M. et al. A collection of breast cancer cell lines for the study of functionally distinct cancer subtypes. Cancer Cell 10, 515–527, doi:10.1016/j.ccr.2006.10.008 (2006).

13 Tsherniak, A. et al. Defining a Cancer Dependency Map. Cell 170, 564–576 e516, doi:10.1016/j.cell.2017.06.010 (2017).

14 Watson, S. S. et al. Microenvironment-Mediated Mechanisms of Resistance to HER2 Inhibitors Differ between HER2+ Breast Cancer Subtypes. Cell Syst 6, 329–342 e326, doi:10.1016/j.cels.2018.02.001 (2018).

15 Wilson, T. R. et al. Widespread potential for growth-factor-driven resistance to anticancer kinase inhibitors. Nature 487, 505–509, doi:10.1038/nature11249 (2012).

16 Morrison, D. K. MAP kinase pathways. Cold Spring Harb Perspect Biol 4, doi:10.1101/cshperspect.a011254 (2012).

17 Harrison, D. A. The Jak/STAT pathway. Cold Spring Harb Perspect Biol 4, doi:10.1101/cshperspect.a011205 (2012).

18 Nusse, R. Wnt signaling. Cold Spring Harb Perspect Biol 4, doi:10.1101/cshperspect.a011163 (2012).

19 David, C. J. & Massague, J. Contextual determinants of TGFbeta action in development, immunity and cancer. Nat Rev Mol Cell Biol 19, 419–435, doi:10.1038/s41580-018-0007-0 (2018).

20 Subramanian, A. et al. A Next Generation Connectivity Map: L1000 Platform and the First 1,000,000 Profiles. Cell 171, 1437–1452 e1417, doi:10.1016/j.cell.2017.10.049 (2017).

21 Zhao, W. et al. Large-Scale Characterization of Drug Responses of Clinically Relevant Proteins in Cancer Cell Lines. Cancer Cell 38, 829–843 e824, doi:10.1016/j.ccell.2020.10.008 (2020).

22 Ng, P. K. et al. Systematic Functional Annotation of Somatic Mutations in Cancer. Cancer Cell 33, 450–462 e410, doi:10.1016/j.ccell.2018.01.021 (2018).

23 Bock, C. et al. The Organoid Cell Atlas. Nat Biotechnol 39, 13–17, doi:10.1038/s41587-020-00762-x (2021).

24 Drost, J. & Clevers, H. Organoids in cancer research. Nat Rev Cancer 18, 407–418, doi:10.1038/s41568-018-0007-6 (2018).

25 Sullivan, L. F. Rewiring the Drosophila Brain With Genetic Manipulations in Neural Lineages. Front Mol Neurosci 12, 82, doi:10.3389/fnmol.2019.00082 (2019).

26 Kinser, H. E. & Pincus, Z. High-throughput screening in the C. elegans nervous system. Mol Cell Neurosci 80, 192–197, doi:10.1016/j.mcn.2016.06.001 (2017).

27 Srinivasan, J. et al. A modular library of small molecule signals regulates social behaviors in Caenorhabditis elegans. PLoS Biol 10, e1001237, doi:10.1371/journal.pbio.1001237 (2012).

28 Saydmohammed, M. & Tsang, M. High-Throughput Automated Chemical Screens in Zebrafish. Methods Mol Biol 1683, 383–393, doi:10.1007/978-1-4939-7357-6_22 (2018).

29 Kersten, K., de Visser, K. E., van Miltenburg, M. H. & Jonkers, J. Genetically engineered mouse models in oncology research and cancer medicine. EMBO Mol Med 9, 137–153, doi:10.15252/emmm.201606857 (2017).

30 Rappoport, N. & Shamir, R. Multi-omic and multi-view clustering algorithms: review and cancer benchmark. Nucleic Acids Res 46, 10546–10562, doi:10.1093/nar/gky889 (2018).

31 Saelens, W., Cannoodt, R. & Saeys, Y. A comprehensive evaluation of module detection methods for gene expression data. Nat Commun 9, 1090, doi:10.1038/s41467-018-03424-4 (2018).

32 Jojic, V. et al. Identification of transcriptional regulators in the mouse immune system. Nat Immunol 14, 633–643, doi:10.1038/ni.2587 (2013).

33 Yosef, N. et al. Dynamic regulatory network controlling TH17 cell differentiation. Nature 496, 461–468, doi:10.1038/nature11981 (2013).

34 Alsina, L. et al. A narrow repertoire of transcriptional modules responsive to pyogenic bacteria is impaired in patients carrying loss-of-function mutations in MYD88 or IRAK4. Nat Immunol 15, 1134–1142, doi:10.1038/ni.3028 (2014).

35 Consortium, E. P. An integrated encyclopedia of DNA elements in the human genome. Nature 489, 57–74, doi:10.1038/nature11247 (2012).

36 Cancer Genome Atlas Research, N. et al. The Cancer Genome Atlas Pan-Cancer analysis project. Nat Genet 45, 1113–1120, doi:10.1038/ng.2764 (2013).

37 Consortium, G. T. The GTEx Consortium atlas of genetic regulatory effects across human tissues. Science 369, 1318–1330, doi:10.1126/science.aaz1776 (2020).

38 Hu, B. C. The human body at cellular resolution: the NIH Human Biomolecular Atlas Program. Nature 574, 187–192, doi:10.1038/s41586-019-1629-x (2019).

39 Soule, H. D. et al. Isolation and characterization of a spontaneously immortalized human breast epithelial cell line, MCF-10. Cancer Res 50, 6075–6086 (1990).

40 Witt, A. E. et al. Functional proteomics approach to investigate the biological activities of cDNAs implicated in breast cancer. J Proteome Res 5, 599–610, doi:10.1021/pr050395r (2006).

41 McQueen, C. M. et al. PER2 regulation of mammary gland development. Development 145, doi:10.1242/dev.157966 (2018).

42 Melani, M., Simpson, K. J., Brugge, J. S. & Montell, D. Regulation of cell adhesion and collective cell migration by hindsight and its human homolog RREB1. Curr Biol 18, 532–537, doi:10.1016/j.cub.2008.03.024 (2008).

43 Seton-Rogers, S. E. et al. Cooperation of the ErbB2 receptor and transforming growth factor beta in induction of migration and invasion in mammary epithelial cells. Proc Natl AcadSci USA 101, 1257–1262, doi:10.1073/pnas.0308090100 (2004).

44 Debnath, J. et al. The role of apoptosis in creating and maintaining luminal space within normal and oncogene-expressing mammary acini. Cell 111, 29–40, doi:10.1016/s0092-8674(02)01001-2 (2002).

45 Debnath, J., Muthuswamy, S. K. & Brugge, J. S. Morphogenesis and oncogenesis of MCF-10A mammary epithelial acini grown in three-dimensional basement membrane cultures. Methods 30, 256–268, doi:10.1016/s1046-2023(03)00032-x (2003).

46 Smith, R. et al. Using Microarrays to Interrogate Microenvironmental Impact on Cellular Phenotypes in Cancer. J Vis Exp, doi:10.3791/58957 (2019).

47 Lin, J. R., Fallahi-Sichani, M., Chen, J. Y. & Sorger, P. K. Cyclic Immunofluorescence (CycIF), A Highly Multiplexed Method for Single-cell Imaging. Curr Protoc Chem Biol 8, 251–264, doi:10.1002/cpch.14 (2016).

48 Lin, J. R., Fallahi-Sichani, M. & Sorger, P. K. Highly multiplexed imaging of single cells using a high-throughput cyclic immunofluorescence method. Nat Commun 6, 8390, doi:10.1038/ncomms9390 (2015).

49 Niepel, M. et al. A Multi-center Study on the Reproducibility of Drug-Response Assays in Mammalian Cell Lines. Cell Syst 9, 35–48 e35, doi:10.1016/j.cels.2019.06.005 (2019).

50 Tibes, R. et al. Reverse phase protein array: validation of a novel proteomic technology and utility for analysis of primary leukemia specimens and hematopoietic stem cells. Mol Cancer Ther 5, 2512–2521, doi:10.1158/1535-7163.MCT-06-0334 (2006).

51 Creech, A. L. et al. Building the Connectivity Map of epigenetics: chromatin profiling by quantitative targeted mass spectrometry. Methods 72, 57–64, doi:10.1016/j.ymeth.2014.10.033 (2015).

52 Abd El-Rehim, D. M. et al. Expression of luminal and basal cytokeratins in human breast carcinoma. J Pathol 203, 661–671, doi:10.1002/path.1559 (2004).

53 McInnes, L., Healy, J. & Melville, J. UMAP: Uniform Manifold Approximation and Projection for Dimension Reduction. arXiv arXiv:1802.03426(2018).

54 Koh, A. S. et al. Rapid chromatin repression by Aire provides precise control of immune tolerance. Nat Immunol 19, 162–172, doi:10.1038/s41590-017-0032-8 (2018).

55 Moskowitz, D. M. & Greenleaf, W. J. Nonparametric analysis of contributions to variance in genomics and epigenomics data. biorxiv, doi:https://doi.org/10.1101/314112 (2018).

56 Subramanian, A. et al. Gene set enrichment analysis: a knowledge-based approach for interpreting genome-wide expression profiles. Proc Natl Acad Sci U S A 102, 15545–15550, doi:10.1073/pnas.0506580102 (2005).

57 Stathias, V. et al. Drug and disease signature integration identifies synergistic combinations in glioblastoma. Nat Commun 9, 5315, doi:10.1038/s41467-018-07659-z (2018).

58 Ivashkiv, L. B. IFNgamma: signalling, epigenetics and roles in immunity, metabolism, disease and cancer immunotherapy. Nat Rev Immunol 18, 545–558, doi:10.1038/s41577-018-0029-z (2018).

59 Belinky, F. et al. PathCards: multi-source consolidation of human biological pathways. Database (Oxford) 2015, doi:10.1093/database/bav006 (2015).

60 Carrasco Pro, S. et al. Global landscape of mouse and human cytokine transcriptional regulation. Nucleic Acids Res 46, 9321–9337, doi:10.1093/nar/gky787 (2018).

61 Mok, S. et al. Inhibition of CSF-1 receptor improves the antitumor efficacy of adoptive cell transfer immunotherapy. Cancer Res 74, 153–161, doi:10.1158/0008-5472.CAN-13-1816 (2014).

62 Zhu, Y. et al. CSF1/CSF1R blockade reprograms tumor-infiltrating macrophages and improves response to T-cell checkpoint immunotherapy in pancreatic cancer models. Cancer Res 74, 5057–5069, doi:10.1158/0008-5472.CAN-13-3723 (2014).

63 Zhao, M. et al. Development of a recombinant human IL-15.sIL-15Ralpha/Fc superagonist with improved half-life and its antitumor activity alone or in combination with PD-1 blockade in mouse model. Biomed Pharmacother 112, 108677, doi:10.1016/j.biopha.2019.108677 (2019).

64 Berraondo, P., Etxeberria, I., Ponz-Sarvise, M. & Melero, I. Revisiting Interleukin-12 as a Cancer Immunotherapy Agent. Clin Cancer Res 24, 2716–2718, doi:10.1158/1078-0432.CCR-18-0381 (2018).

65 Flores-Toro, J. A. et al. CCR2 inhibition reduces tumor myeloid cells and unmasks a checkpoint inhibitor effect to slow progression of resistant murine gliomas. Proc Natl Acad Sci U S A 117, 1129–1138, doi:10.1073/pnas.1910856117 (2020).

66 Steele, C. W. et al. CXCR2 Inhibition Profoundly Suppresses Metastases and Augments Immunotherapy in Pancreatic Ductal Adenocarcinoma. Cancer Cell 29, 832–845, doi:10.1016/j.ccell.2016.04.014 (2016).

67 Hunt, G. J., Dane, M. A., Korkola, J. E., Heiser, L. M. & Gagnon-Bartsch, J. A. Automatic Transformation and Integration to Improve Visualization and Discovery of Latent Effects in Imaging Data. Journal of Computational and Graphical Statistics 29, 929–941 (2019).

68 Keenan, A. B. et al. ChEA3: transcription factor enrichment analysis by orthogonal omics integration. Nucleic Acids Res 47, W212–W224, doi:10.1093/nar/gkz446 (2019).

69 Yirmiya, R. & Goshen, I. Immune modulation of learning, memory, neural plasticity and neurogenesis. Brain Behav Immun 25, 181–213, doi:10.1016/j.bbi.2010.10.015 (2011).

70 Good, A. L. & Stoffers, D. A. Stress-Induced Translational Regulation Mediated by RNA Binding Proteins: Key Links to beta-Cell Failure in Diabetes. Diabetes 69, 499–507, doi:10.2337/dbi18-0068 (2020).

71 Moss, N. D. & Sussel, L. mRNA Processing: An Emerging Frontier in the Regulation of Pancreatic beta Cell Function. Front Genet 11, 983, doi:10.3389/fgene.2020.00983 (2020).

72 Wagenseil, J. E. & Mecham, R. P. Vascular extracellular matrix and arterial mechanics. Physiol Rev 89, 957–989, doi:10.1152/physrev.00041.2008 (2009).

73 Witjas, F. M. R., van den Berg, B. M., van den Berg, C. W., Engelse, M. A. & Rabelink, T. J. Concise Review: The Endothelial Cell Extracellular Matrix Regulates Tissue Homeostasis and Repair. Stem Cells Transl Med 8, 375–382, doi:10.1002/sctm.18-0155 (2019).

74 Heng, B. C. et al. Role of YAP/TAZ in Cell Lineage Fate Determination and Related Signaling Pathways. Front Cell Dev Biol 8, 735, doi:10.3389/fcell.2020.00735 (2020).

75 Cohen, D. R. & Curran, T. fra-1: a serum-inducible, cellular immediate-early gene that encodes a fos-related antigen. Mol Cell Biol 8, 2063–2069, doi:10.1128/mcb.8.5.2063-2069.1988 (1988).

76 Cohen, D. R., Ferreira, P. C., Gentz, R., Franza, B. R., Jr. & Curran, T. The product of a fos-related gene, fra-1, binds cooperatively to the AP-1 site with Jun: transcription factor AP-1 is comprised of multiple protein complexes. Genes Dev 3, 173–184, doi:10.1101/gad.3.2.173 (1989).

77 Gillies, T. E., Pargett, M., Minguet, M., Davies, A. E. & Albeck, J. G. Linear Integration of ERK Activity Predominates over Persistence Detection in Fra-1 Regulation. Cell Syst 5, 549–563 e545, doi:10.1016/j.cels.2017.10.019 (2017).

78 Rutter, J., Michnoff, C. H., Harper, S. M., Gardner, K. H. & McKnight, S. L. PAS kinase: an evolutionarily conserved PAS domain-regulated serine/threonine kinase. Proc Natl Acad Sci U S A 98, 8991–8996, doi:10.1073/pnas.161284798 (2001).

79 Wilson, W. A. et al. Control of mammalian glycogen synthase by PAS kinase. Proc Natl AcadSci USA 102, 16596–16601, doi:10.1073/pnas.0508481102 (2005).

80 Tirosh, I. et al. Dissecting the multicellular ecosystem of metastatic melanoma by singlecell RNA-seq. Science 352, 189–196, doi:10.1126/science.aad0501 (2016).

81 Hafner, M. et al. Multiomics Profiling Establishes the Polypharmacology of FDA-Approved CDK4/6 Inhibitors and the Potential for Differential Clinical Activity. Cell Chem Biol 26, 1067–1080 e1068, doi:10.1016/j.chembiol.2019.05.005 (2019).

82 Jones, S. A. & Jenkins, B. J. Recent insights into targeting the IL-6 cytokine family in inflammatory diseases and cancer. Nat Rev Immunol 18, 773–789, doi:10.1038/s41577-018-0066-7 (2018).

83 Ng, M. R., Besser, A., Danuser, G. & Brugge, J. S. Substrate stiffness regulates cadherin-dependent collective migration through myosin-II contractility. J Cell Biol 199, 545–563, doi:10.1083/jcb.201207148 (2012).

84 Poplimont, H. et al. Neutrophil Swarming in Damaged Tissue Is Orchestrated by Connexins and Cooperative Calcium Alarm Signals. Curr Biol 30, 2761–2776 e2767, doi:10.1016/j.cub.2020.05.030 (2020).

85 Huang, R. et al. The NCATS BioPlanet - An Integrated Platform for Exploring the Universe of Cellular Signaling Pathways for Toxicology, Systems Biology, and Chemical Genomics. Front Pharmacol 10, 445, doi:10.3389/fphar.2019.00445 (2019).

86 Amara, U. et al. Molecular intercommunication between the complement and coagulation systems. J Immunol 185, 5628–5636, doi:10.4049/jimmunol.0903678 (2010).

87 Bezdek, J. C., Ehrlich, R. & Full, W. FCM: The fuzzy c-means clustering algorithm. Computers & Geosciences 10(1984).

88 Bonnans, C., Chou, J. & Werb, Z. Remodelling the extracellular matrix in development and disease. Nat Rev Mol Cell Biol 15, 786–801, doi:10.1038/nrm3904 (2014).

89 Magnusson, K. E., Jalden, J., Gilbert, P. M. & Blau, H. M. Global linking of cell tracks using the Viterbi algorithm. IEEE Trans Med Imaging 34, 911–929, doi:10.1109/TMI.2014.2370951 (2015).

90 Schindelin, J. et al. Fiji: an open-source platform for biological-image analysis. Nat Methods 9, 676–682, doi:10.1038/nmeth.2019 (2012).

91 Meijering, E., Dzyubachyk, O. & Smal, I. Chapter nine - Methods for Cell and Particle Tracking. Methods in Enzymology (2012).

92 Berg, S. et al. ilastik: interactive machine learning for (bio)image analysis. Nat Methods 16, 1226–1232, doi:10.1038/s41592-019-0582-9 (2019).

93 McQuin, C. et al. CellProfiler 3.0: Next-generation image processing for biology. PLoS Biol 16, e2005970, doi:10.1371/journal.pbio.2005970 (2018).

94 Ebdon, D. Statistics in geography. (1985).

95 Akbani, R. et al. A pan-cancer proteomic perspective on The Cancer Genome Atlas. Nat Commun 5, 3887, doi:10.1038/ncomms4887 (2014).

96 Tatlow, P. J. & Piccolo, S. R. A cloud-based workflow to quantify transcript-expression levels in public cancer compendia. Sci Rep 6, 39259, doi:10.1038/srep39259 (2016).

97 Love, M. I., Huber, W. & Anders, S. Moderated estimation of fold change and dispersion for RNA-seq data with DESeq2. Genome Biol 15, 550, doi:10.1186/s13059-014-0550-8 (2014).

98 Hanzelmann, S., Castelo, R. & Guinney, J. GSVA: gene set variation analysis for microarray and RNA-seq data. BMC Bioinformatics 14, 7, doi:10.1186/1471-2105-14-7 (2013).

99 Corces, M. R. et al. An improved ATAC-seq protocol reduces background and enables interrogation of frozen tissues. Nat Methods 14, 959–962, doi:10.1038/nmeth.4396 (2017).

100 Ross-Innes, C. S. et al. Differential oestrogen receptor binding is associated with clinical outcome in breast cancer. Nature 481, 389–393, doi:10.1038/nature10730 (2012).

101 Schep, A. N., Wu, B., Buenrostro, J. D. & Greenleaf, W. J. chromVAR: inferring transcription-factor-associated accessibility from single-cell epigenomic data. Nat Methods 14, 975–978, doi:10.1038/nmeth.4401 (2017).

102 Khan, A. et al. JASPAR 2018: update of the open-access database of transcription factor binding profiles and its web framework. Nucleic Acids Res 46, D260–D266, doi:10.1093/nar/gkx1126 (2018).

103 Peng, T. et al. A BaSiC tool for background and shading correction of optical microscopy images. Nat Commun 8, 14836, doi:10.1038/ncomms14836 (2017).

104 Wang, Z., Lachmann, A., Keenan, A. B. & Ma’ayan, A. L1000FWD: fireworks visualization of drug-induced transcriptomic signatures. Bioinformatics 34, 2150–2152, doi:10.1093/bioinformatics/bty060 (2018).

105 Wickham, H. et al. Welcome to the Tidyverse. The Journal of Open Source Software 4(43)(2019).

