## Supplementary Figures for "A LINCS microenvironment perturbation resource for integrative assessment of ligand-mediated molecular and phenotypic responses"

#### Sup. Fig. 1 Experimental and bioinformatic approaches to identify high impact ligands

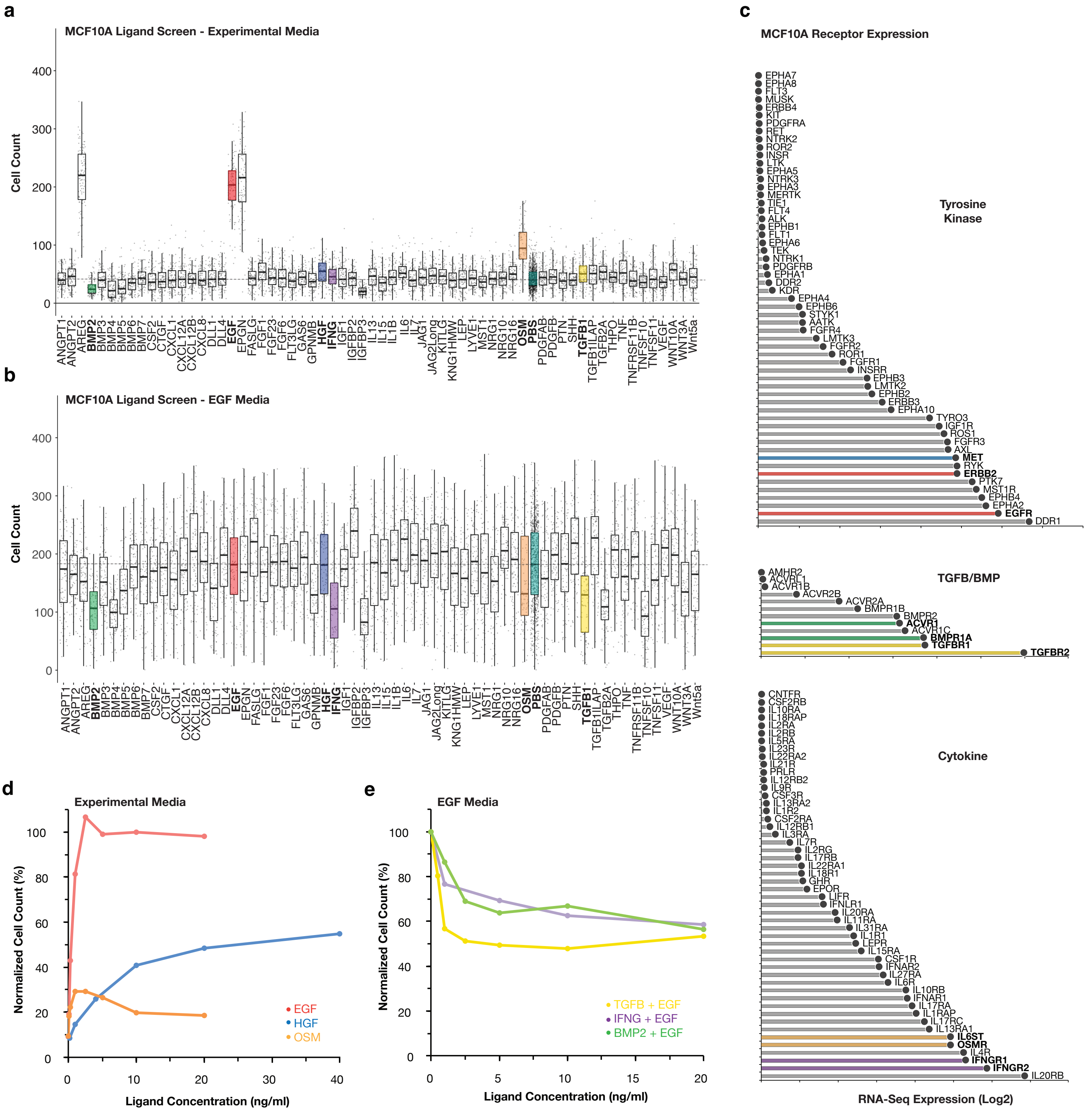

**Sup. Fig. 2 Comparison of ligand and small molecule inhibitor signatures**

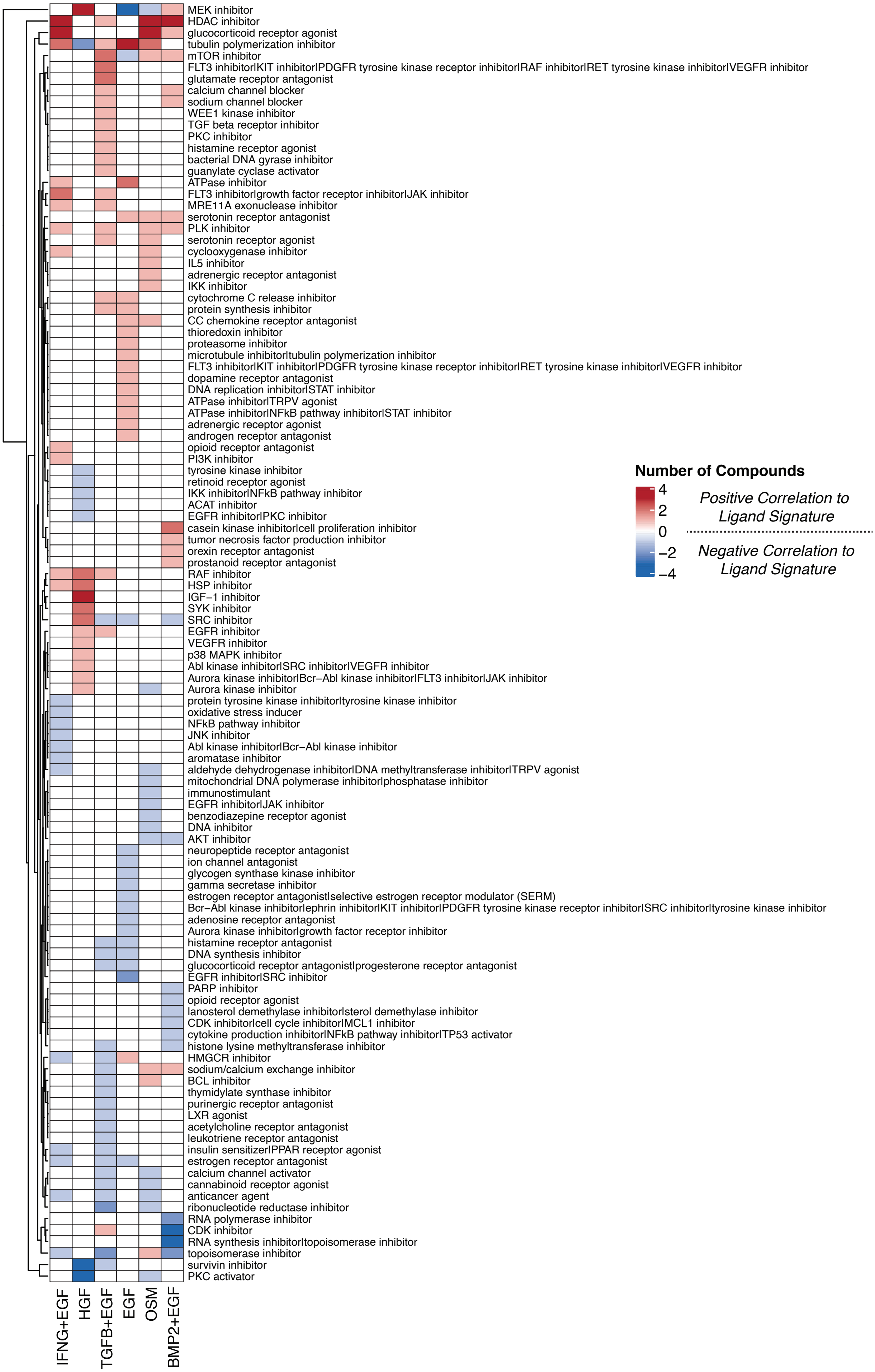

**Sup. Fig. 3 IFNG responses are dynamically encoded across multiple molecular modalities**

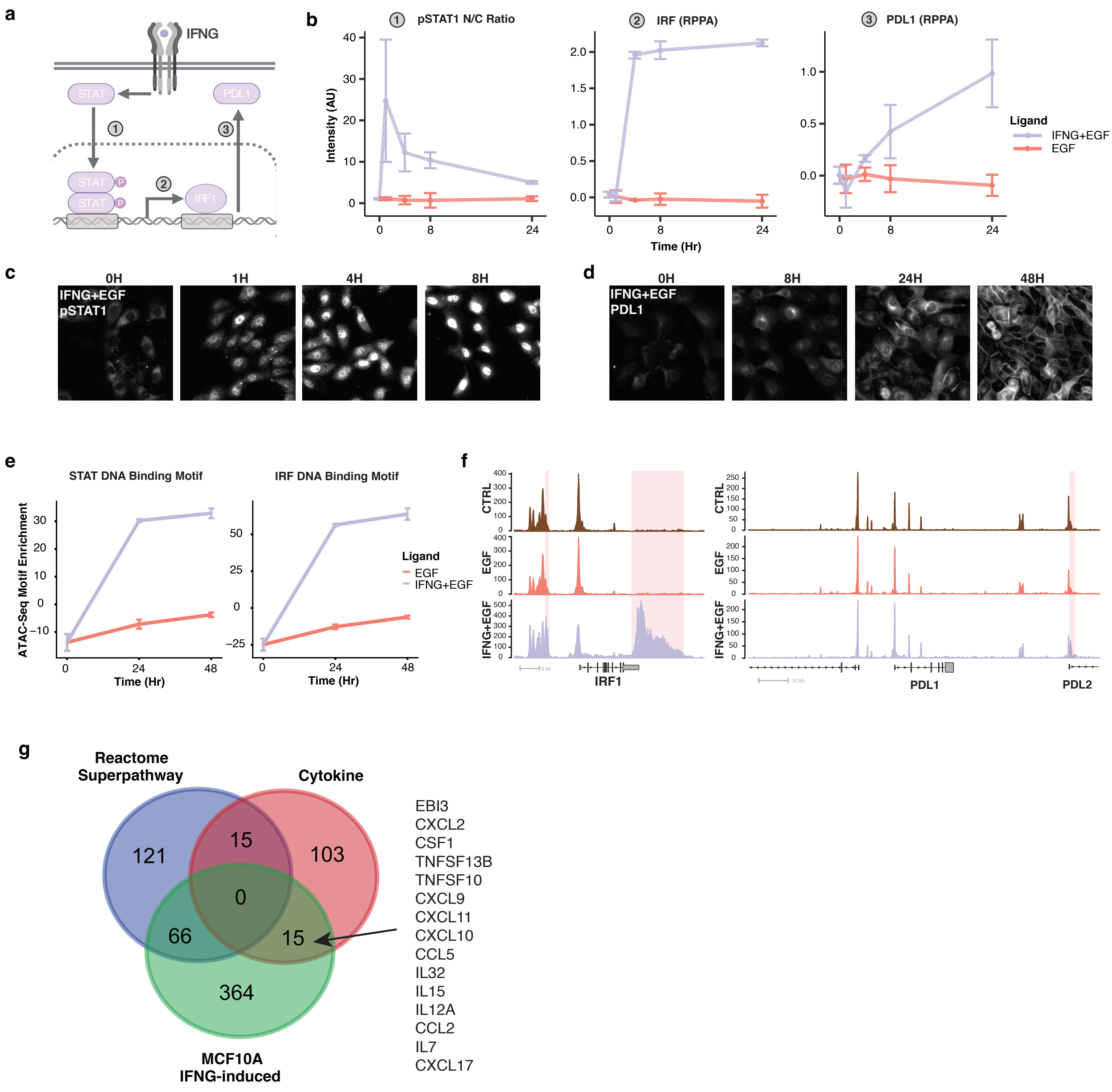

**Sup. Fig. 4 Comparison across RPPA, RNAseq and ATACseq assays reveals concordance across molecular modalities in response to ligand treatment**

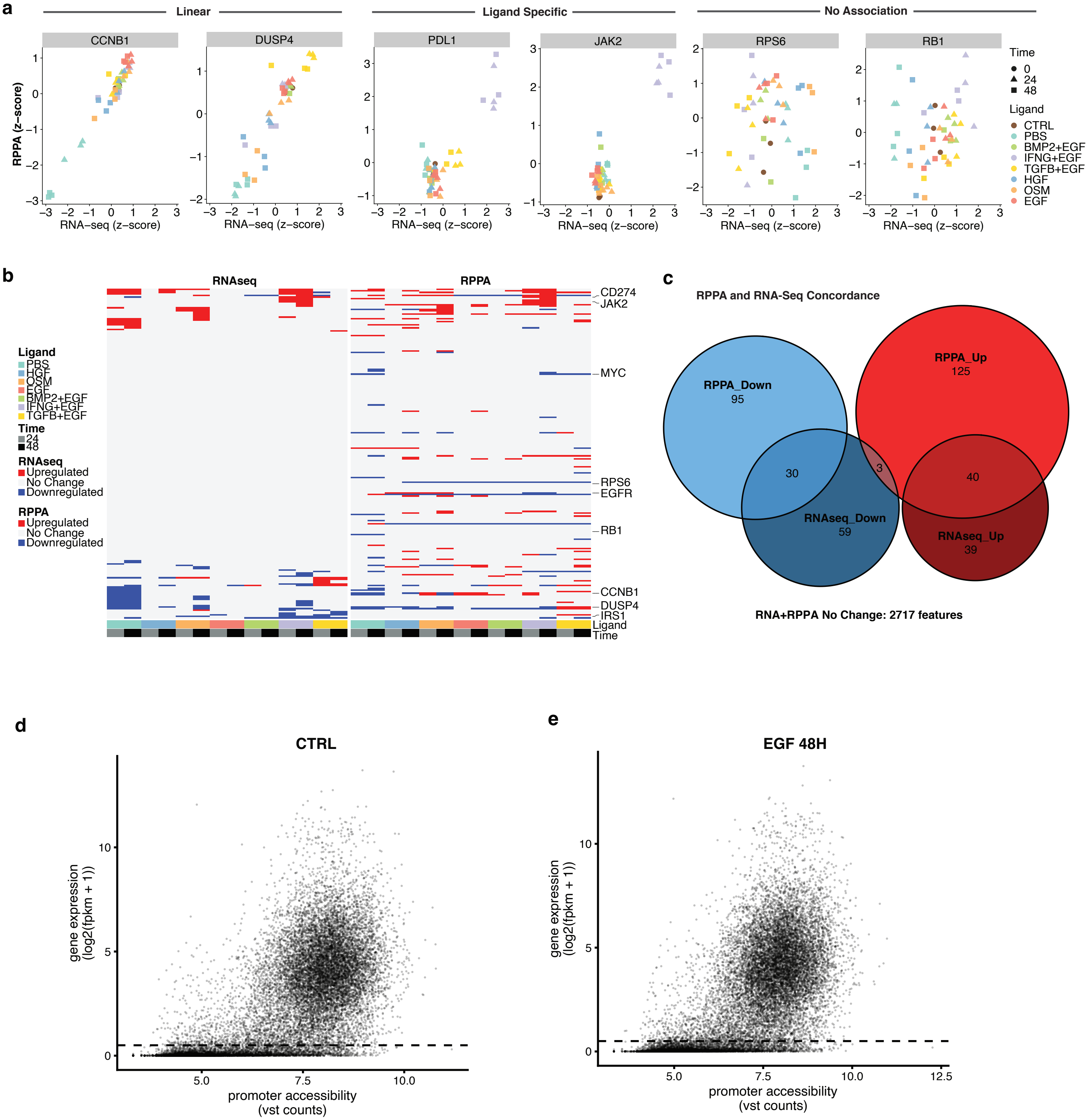

### MDD Integrated Analysis

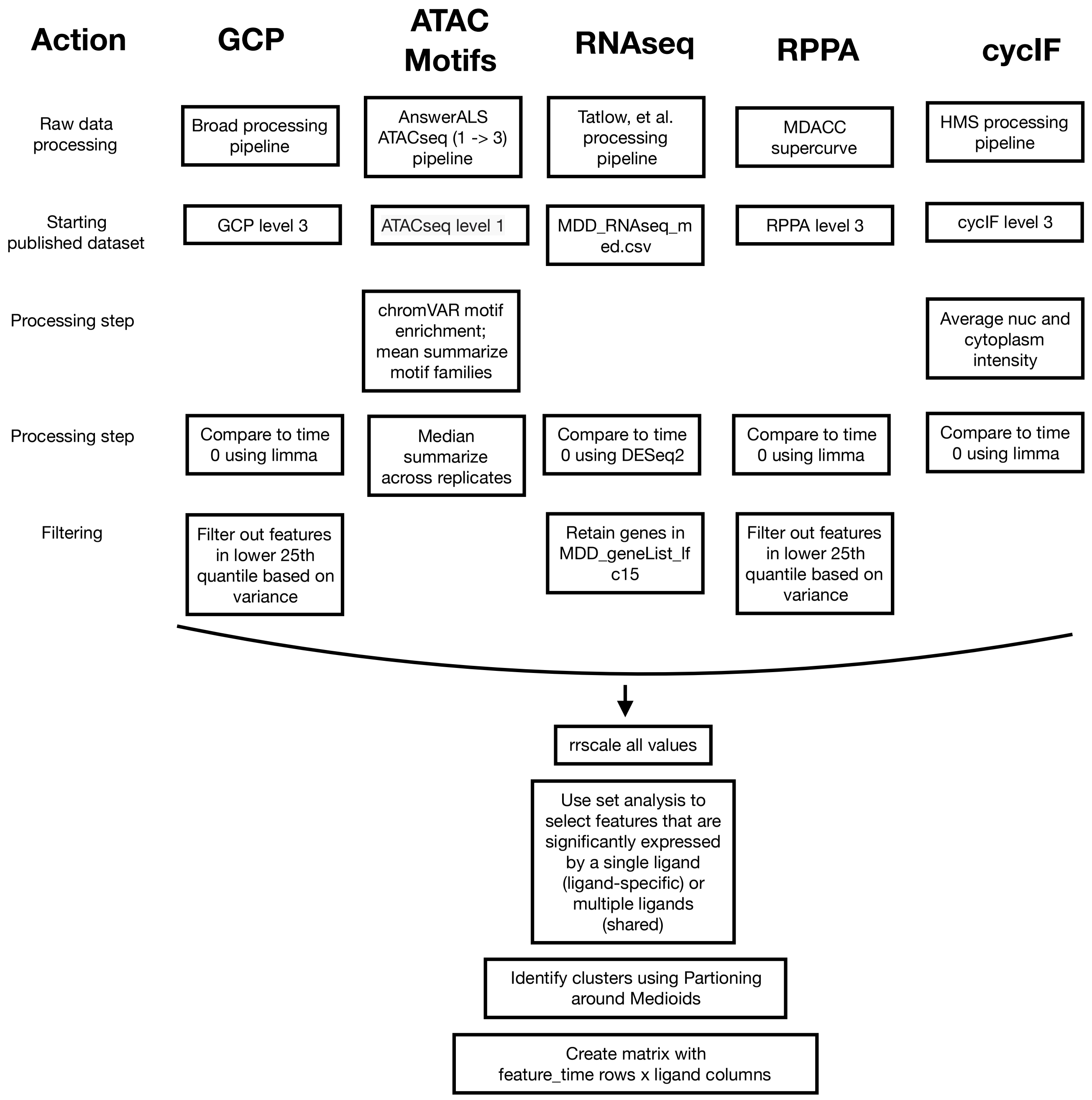

**Sup. Fig. 6 Identification and characterization of integrative molecular modules**

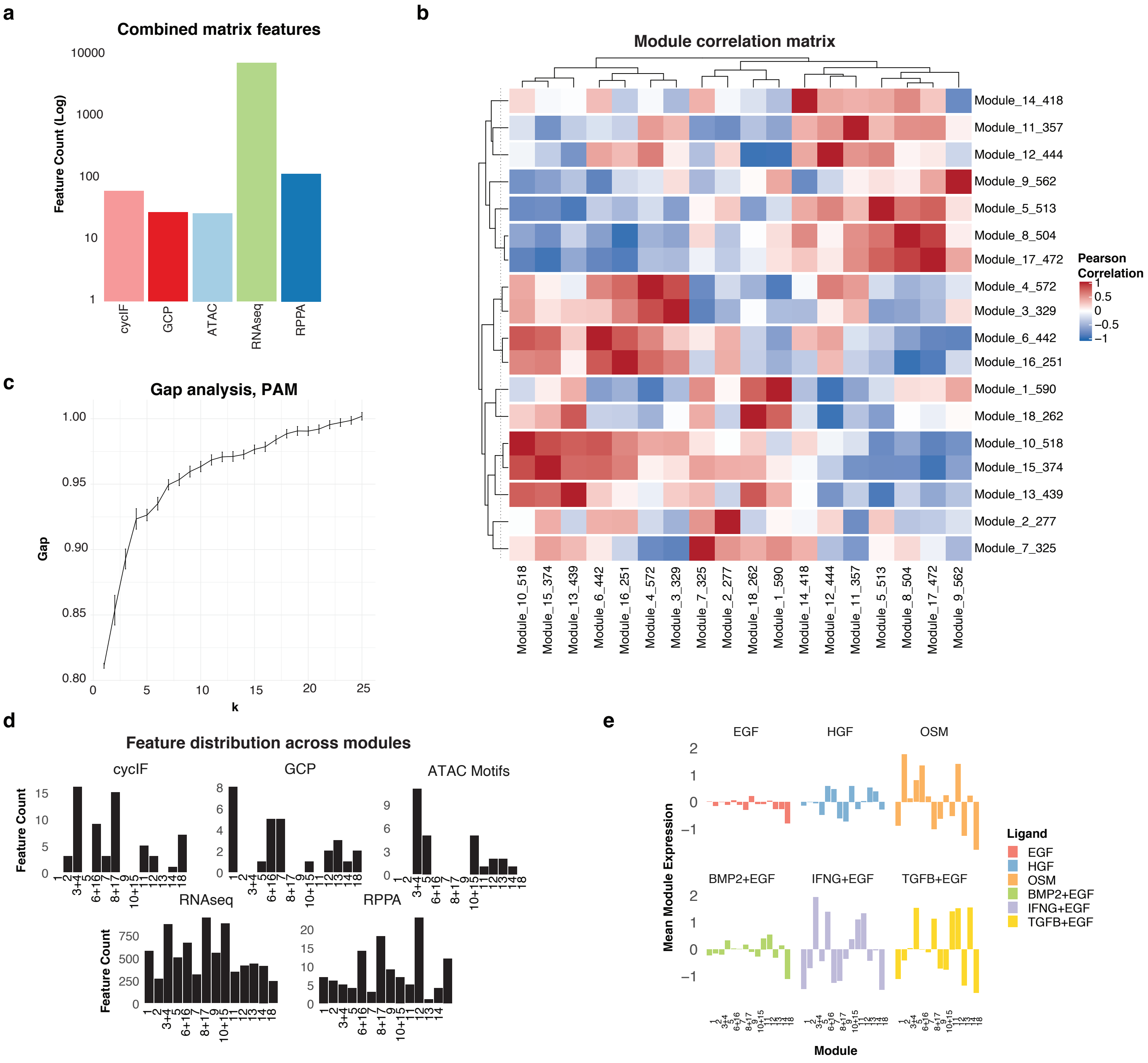

**Sup. Fig. 7 GTEx expression analysis**

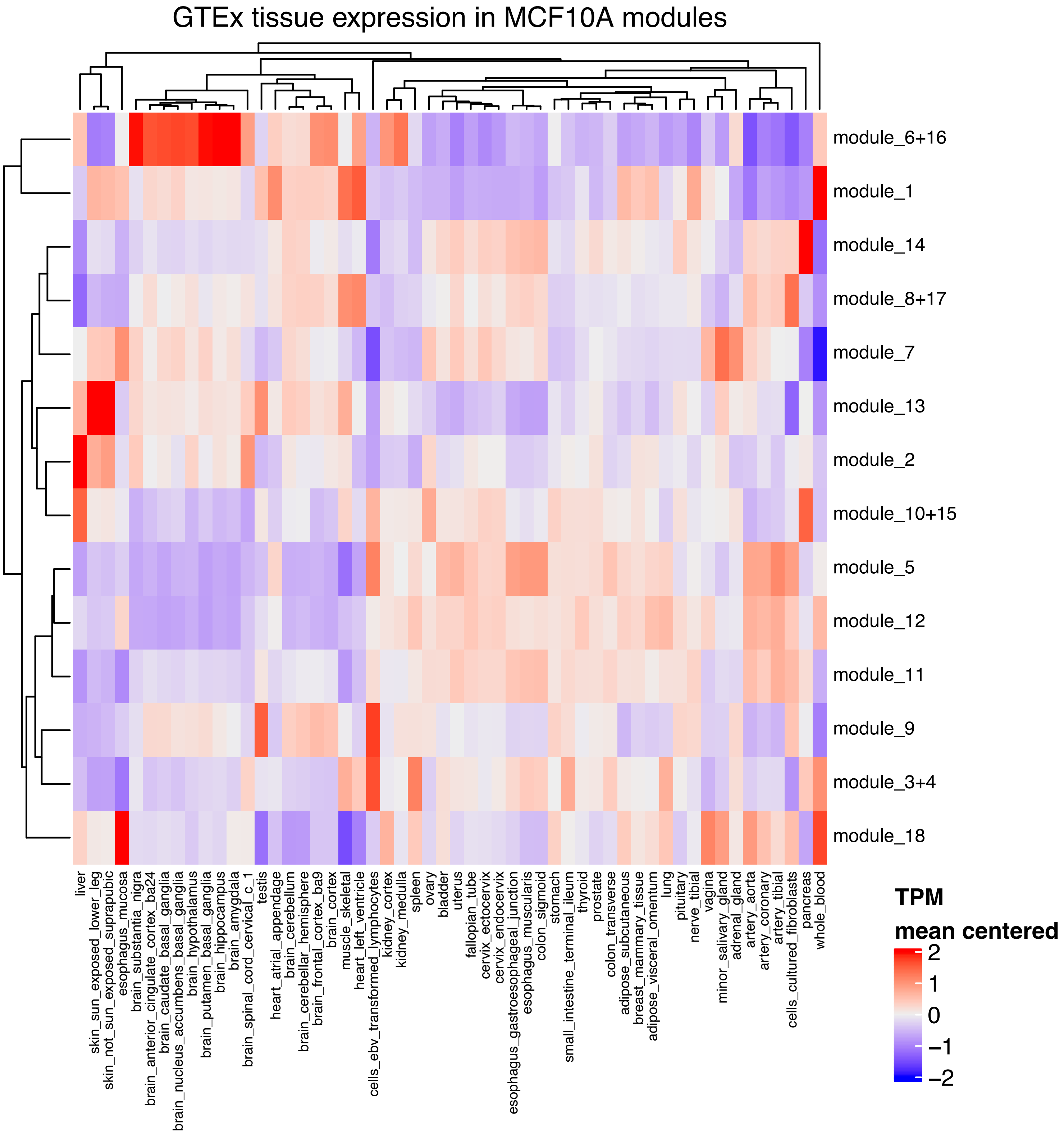
